# The DNA demethylase TET1 recognises chromatin via nucleosome engagement of its N-terminal region

**DOI:** 10.64898/2026.10.31.763349

**Authors:** Franck Martin, Hayden Burdett, Mélody Matelot, Chloé Ngon Baroung, Beatrice Alexander-Howden, Marcus D. Wilson, Elisa Bergamin, Raphaël Pantier

**Author notes:** These authors contributed equally (Franck Martin, Hayden Burdett).

## Abstract

TET1 is a DNA demethylase critical for mammalian embryonic development, yet how it is recruited to its chromatin substrate remains poorly understood. Here we identify a previously uncharacterized N-terminal region of TET1 that directly binds nucleosomes and promotes chromatin binding in mouse embryonic stem cells. Robust chromatin association was evidenced by fluorescence recovery after photobleaching during interphase and chromosome retention throughout mitosis. A 2.47 Å cryo-EM structure of the TET1-nucleosome complex reveals recognition of the nucleosome acidic patch through a canonical arginine anchor centred on R75, with additional contacts extending along the H2B surface. Biochemical and cellular analyses validate this interaction and demonstrate its contribution to TET1 chromatin engagement. The region is absent from TET2 and TET3 and displays strong conservation within mammals, indicating that direct nucleosome recognition is a specialized feature of TET1. Together, these findings uncover a previously unknown chromatin-binding module in TET1 and establish a structural basis for its interaction with nucleosomes.

## Introduction

Ten-eleven translocation (TET) proteins are important epigenetic regulators responsible for the oxidation of 5-methylcytosine into 5-hydroxymethylcytosine, 5-formylcytosine and 5-carboxylcytosine, thereby contributing to active DNA demethylation of the genome ^1–4^. In mammals, the TET family is comprised of three paralogue proteins (TET1, TET2, TET3) sharing a conserved C-terminal catalytic domain. TET function is critical for a number of physiological processes, including embryonic development, haematopoiesis, gametogenesis, immune reaction and brain activity ^5,6^. Conversely, dysregulation of TET proteins is responsible for cancers and developmental disorders ^7–9^. Genetic deletion studies in stem cells and animal models have demonstrated that TET proteins act synergistically to control early cell fate decisions during embryonic development ^10–15^. However, deletion of single paralogues is also known to cause defects in specific organs or developmental processes ^16–20^. In particular, TET1 is the most prominently expressed TET paralogue in pluripotent stem cells ^21,22^. *Tet1* knockout affects lineage specification in cellular models ^23–26^, and results in embryonic lethality in transgenic mouse models with variable penetrance depending on genetic background ^16,19,23^.

The molecular basis underlying the unique and redundant functions of TET paralogues remains incompletely understood. TET protein paralogues exhibit tissue-specific expression patterns, in part explaining their distinct biological activity ^5,27,28^. Additionally, while the C-terminal catalytic domain (conferring DNA demethylase activity) is a highly conserved feature, TET proteins carry a large N-terminal region (∼1,000 amino acids) ^4^ which is considerably divergent between paralogues. These N-terminal regions remain poorly characterised; however, the conservation of some segments across species suggests they could potentially contain important functional domains. The only known chromatin-binding domain within TET proteins is a CXXC zinc-finger domain ^29^. Unlike most CXXC domains, the TET CXXC domain exhibits broad sequence specificity, recognising an unmethylated cytosine followed by diverse nucleotides, including 5-methylcytosine-containing sequences ^29–32^. Despite its biological importance ^30,33^, the CXXC domain cannot fully explain specific genomic targeting, in particular over enhancer and promoter regulatory elements which are actively demethylated by TETs ^34–40^. Indeed, TET2 has lost its ancestral CXXC domain entirely ^41^ and several isoforms of TET1/3 lack the CXXC but retain chromatin binding activity ^33,42–44^, showing the dispensability of this domain in some cases. These observations indicate that the CXXC domain is dispensable in certain contexts and strongly support the existence of CXXC-independent recruitment mechanisms. Such mechanisms likely involve interactions with additional chromatin-associated factors, including sequence-specific transcription factors ^45–49^.

Here, we uncovered a new chromatin-binding region within TET1 using a combination of live-cell imaging, biochemical and structural approaches. We tested a library of truncations which mapped this functional region to the N-terminal 93 amino acids of the protein. Interestingly, protein alignments and phylogenetic analysis showed that this region is unique to the TET1 paralogue (i.e. absent in TET2/3) and restricted to mammalian species. Microscopy analyses indicated that TET1 N-terminal region acts cooperatively with the CXXC domain for general chromatin binding in stem cells, but is specifically responsible for attachment to condensed chromosomes throughout mitosis. We demonstrate that this region interacts directly with nucleosomes and obtained a cryo-EM structure revealing key interactions with the nucleosome acidic patch. Overall, this study provides new understanding for the interaction between an important epigenetic regulator and chromatin.

## Results

### TET1 is specifically retained on chromatin during mitosis in mouse embryonic stem cells

TET1 and TET2 are highly expressed in mouse embryonic stem cells (ESCs), while TET3 protein is undetectable ^21,22^. However, little is known about the sub-nuclear localisation of individual TET paralogues. To address this question, we used previously generated ESC lines in which endogenous *Tet1* and *Tet2* alleles were tagged with 3x Flag epitope tags ^22^ (*Tet1^Flag^* ESCs and *Tet2^Flag^* ESCs, Fig.1A). Immunofluorescence analysis revealed that both TET1 and TET2 were diffusely distributed throughout the nucleoplasm in the vast majority of cells during interphase. Strikingly, TET1 remained associated with condensed chromosomes in mitotic cells, which are easily distinguishable by microscopy due to their characteristic chromatin compaction in metaphase (Fig.1A). In contrast, TET2 appeared to be excluded from mitotic chromatin (Fig.1A), consistent with the widespread displacement of nucleoplasmic proteins during mitosis. Quantitative analysis of immunofluorescence data confirmed robust co-localisation of TET1 with mitotic chromatin in the vast majority of ESCs (96.7%), whereas TET2 showed almost no overlap (3.6% of only partial co-localisation) (Fig.1B). We further verified TET1 retention throughout all stages of mitosis by co-immunofluorescence with an antibody recognising phosphorylated Histone H3 (Ser10), a marker of mitotic chromatin ^50^ (Fig.1C). Of note, the small fraction of mitotic cells lacking chromatin-associated TET1 appeared to be negative for TET1 expression, rather than exhibiting TET1 exclusion from chromosomes. These data indicate that TET1, but not TET2, can bind to condensed chromatin during mitosis (Fig.1B).

**Figure 1.**
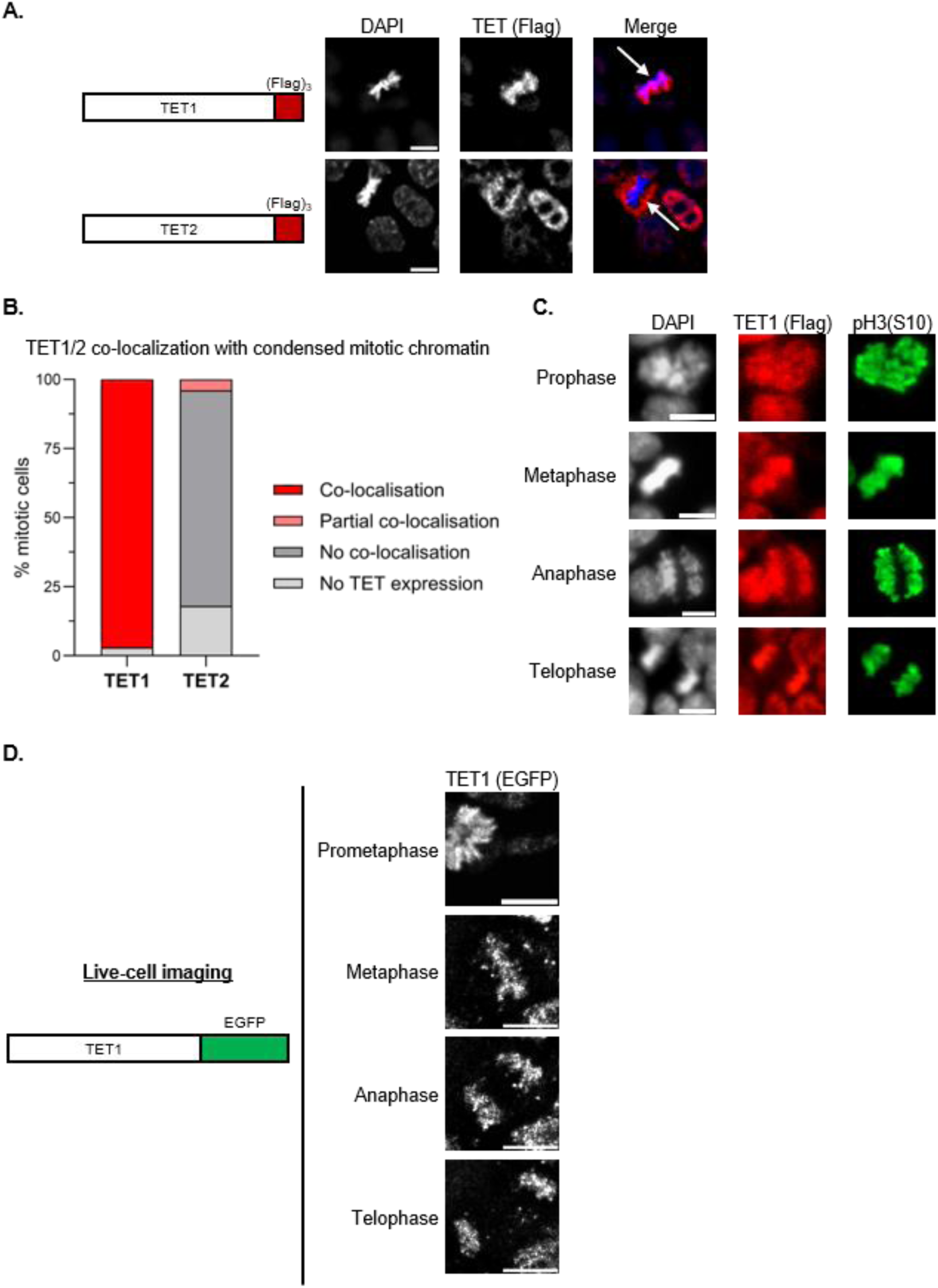
TET1 is specifically associated with chromatin during mitosis. **A**. Immunofluorescence of endogenous TET1 and TET2 tagged with (Flag)_3_ in the indicated mouse embryonic stem cell lines. DNA was stained with DAPI. Arrows indicate mitotic cells. Scale bars: 10µm. **B.** Quantification of microscopy data shown in panel A, focusing on the co-localisation of TET1/2 with condensed mitotic chromatin. **C.** Co-immunofluorescence of endogenous TET1-(Flag)_3_ with phospho-histone H3 (pH3(S10)) at different stages of mitosis in mouse embryonic stem cells. DNA was stained with DAPI. Scale bars: 10µm. **D.** Live-cell imaging of endogenous TET1 tagged with EGFP at different stages of mitosis in mouse embryonic stem cells. Scale bars: 10µm.

As these initial results were obtained from fixed samples, we sought to exclude the possibility that formaldehyde fixation caused artefactual mis-localisation of proteins ^51^. Indeed, proteins which are well known to associate with mitotic chromatin *in vivo*, such as the stem cell factors OCT4 and ESRRB, appear to be completely mis-localised following fixation and immunofluorescence analysis ^52,53^ (Fig.S1A). As an alternative strategy, we performed live-cell imaging in ESCs transiently expressing EGFP-tagged TET1 or TET2. In agreement with immunostaining results, TET1 remains associated with mitotic chromatin in live cells, whereas TET2 is completely excluded from condensed chromosomes (Fig.S1B). These findings demonstrate mitotic chromatin association of TET1 *in vivo*, although transient overexpression could potentially influence its behaviour. To examine TET1 localisation under endogenous conditions, we used CRISPR/Cas9 genome editing in order to tag *Tet1* alleles with a C-terminal EGFP fluorescent reporter (Fig.S1C, S1D). Live-cell imaging of *Tet1^EGFP^* ESCs confirmed that endogenous TET1 remains associated with chromatin throughout mitosis in unperturbed conditions ( Fig.1D, Supplementary video). Together, these results establish that mitotic chromatin retention is an intrinsic property of TET1.

### Mapping a mammalian-specific chromatin-binding region within TET1

The observation that TET1, but not TET2, remains associated with mitotic chromatin prompted us to identify the region responsible for this activity. To this end, we cloned a series of plasmid constructs encoding for EGFP-tagged TET1 truncations, and assessed the localisation of synthesised proteins by live-cell imaging in ESCs (Fig.S2A, S2B). Initial deletion analyses revealed that mitotic chromatin binding was restricted to the N-terminal portion of TET1, whereas the isolated catalytic domain (residues 1367-2039) was unable to associate with mitotic chromosomes (Fig.S2A, S2B). Unexpectedly, TET1 N-terminal fragments lacking the canonical CXXC DNA-binding domain (residues 567-608) still displayed robust chromatin association during mitosis (Fig.2A, 2B), indicating the existence of an additional chromatin-binding module. To further define this activity, we generated a series of progressively truncated N-terminal constructs (Fig.2A). Fragments encompassing residues 1-108 or 1-93 efficiently co-localised with mitotic chromatin, whereas additional deletions from either terminus markedly reduced chromatin association (Fig.2B). Notably, the 52-108 fragment accumulated predominantly in the cytoplasm during interphase (Fig.2B), suggesting that residues within the proximal N-terminal region contribute to nuclear localization. Based on these findings, we defined residues 1-93 as the minimal chromatin-binding region of TET1 and termed this segment the N-terminal (NT) region. We next asked whether the NT region is required for chromatin association in the context of the full-length TET1 protein. To test this hypothesis, we cloned a TET1 mutant lacking residues 1-93 (TET1ΔNT; Fig.2C). Despite retaining predominant nuclear localisation, indicating the presence of additional nuclear localisation determinants elsewhere in the protein, TET1ΔNT displayed a pronounced defect in mitotic chromatin association compared with wild-type TET1 (Fig.2D). Weak residual chromatin binding remained detectable for TET1ΔNT (Fig.2D), likely reflecting the contribution of the intact CXXC domain within this construct. Together, these investigations uncovered a previously unannotated N-terminal region of TET1 that is both necessary and sufficient for robust mitotic chromatin association.

**Figure 2.**
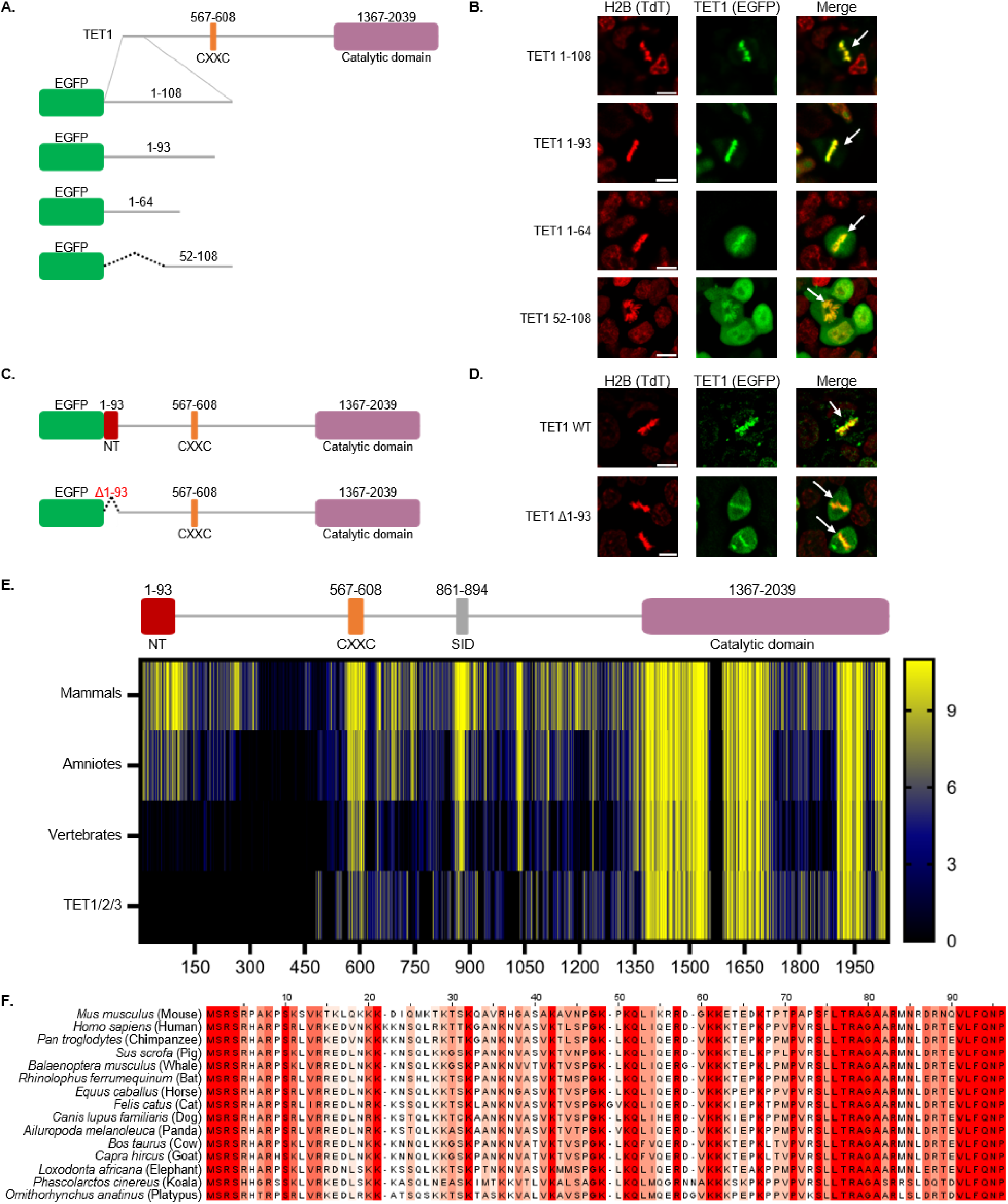
Mapping of an evolutionarily conserved chromatin binding region within TET1 protein. **A.** Diagram showing TET1 protein structure and different truncations cloned into expression plasmids (EGFP fluorescent reporter) to map the region responsible for mitotic chromatin binding. **B.** Live-cell imaging showing the sub-nuclear localisation of the indicated EGFP-TET1 constructs in mouse embryonic stem cells. Chromatin was visualised with Histone H2B fused with a tdTomato fluorescent reporter. Arrows indicate mitotic cells. Scale bars: 10µm. **C.** Diagram showing TET1 wild-type and mutant (Δ1-93) constructs cloned into expression plasmids (EGFP fluorescent reporter) to assess the importance of the N-terminal region (NT) for mitotic chromatin binding. **D.** Live-cell imaging showing the sub-nuclear localisation of the indicated EGFP-TET1 constructs in mouse embryonic stem cells. Chromatin was visualised with Histone H2B fused with a tdTomato fluorescent reporter. Arrows indicate mitotic cells. Scale bars: 10µm.**E.** Heatmap showing the evolutionary conservation of TET1 functional domains across species (mammals, amniotes, vertebrates) and among paralogues (TET1/2/3) within the same gene family. **F.** Protein alignment showing the conservation TET1 N-terminal region (NT) across mammalian species. Strictly conserved residues are highlighted in dark red and conservative substitutions in light red.

Having identified this chromatin-binding region, we next examined its evolutionary conservation. First, protein sequence comparisons across TET paralogues from mouse and human revealed that the NT region is unique to TET1 and absent from both TET2 and TET3 (Fig.S2C). This finding is consistent with the extensive divergence of TET paralogues outside the conserved catalytic domain ^1^, and also with our experimental observation that TET2 is unable to associate with mitotic chromatin (Fig.1A, S1B). Remarkably, the NT region is highly conserved among mammalian TET1 orthologues (Fig.2E, 2F) and is enriched in positively charged residues, a feature consistent with DNA– or chromatin-binding activity. In contrast, conservation of this region was substantially weaker in non-mammalian amniotes (reptiles, birds) and could not be detected in more distantly related vertebrates (Fig.2E, S2D). Notably, other N-terminal TET1 domains, including the CXXC zinc finger ^29^ and Sin3-interacting domain (SID) ^54^, are instead conserved across all vertebrate species (Fig.2E). Collectively, these analyses identify the NT region as a TET1 –specific, mammalian-restricted domain, suggesting that it represents a relatively recent evolutionary innovation.

### The TET1 N-terminal region acts cooperatively with the CXXC to promote TET1 chromatin binding

Our characterisation of the N-terminal chromatin binding region of TET1 was so far limited to its retention on chromatin during mitosis, which is a transient phase of the cell cycle represented by only a small sub-population of cells in culture (<10%). However, we reasoned that this behaviour was unlikely to be specific to mitosis, as many chromatin-associated factors display persistent binding to mitotic chromosomes despite having no established role in cell division ^52,55,56^. We therefore decided to assess whether the NT region also contributes to TET1 chromatin association during interphase. To address this question, we transiently transfected ESCs with plasmids encoding for different TET1 constructs fused with an EGFP tag, and performed fluorescence recovery after photobleaching (FRAP) analysis. We compared the binding dynamics of wild-type TET1 with other variants carrying targeted deletions of the NT region (residues 1-93) and the CXXC domain (residues 567-608), individually or in combination (Fig.3A). As expected, mutation of the well-characterised DNA binding CXXC domain ^29–33^ caused a significant loss in chromatin association characterised by both a shorter half maximum recovery time (t1/2 ∼13sec for TET1 ΔCXXC versus ∼26sec for wild-type TET1) and a higher mobile fraction (plateau ∼93% for TET1 ΔCXXC versus ∼78% for wild-type TET1) (Fig.3B, 3C). Interestingly, the deletion of the NT resulted in a similar quantitative defect (t1/2 ∼18sec and plateau ∼93% for TET1 Δ1-93) as observed for the CXXC deletion (Fig.3B, 3C). Furthermore, combined deletion of these two functional domains caused an even greater loss of chromatin binding activity (t1/2 ∼8sec and plateau ∼99% for TET1 Δ1-93 ΔCXXC) (Fig.3B, 3C). Together, these data demonstrate the NT region has major impact on TET1 chromatin binding activity *in vivo*. The additive effects of NT and CXXC deletions further suggest that these two domains act cooperatively to stabilise TET1 association with chromatin.

**Figure 3.**
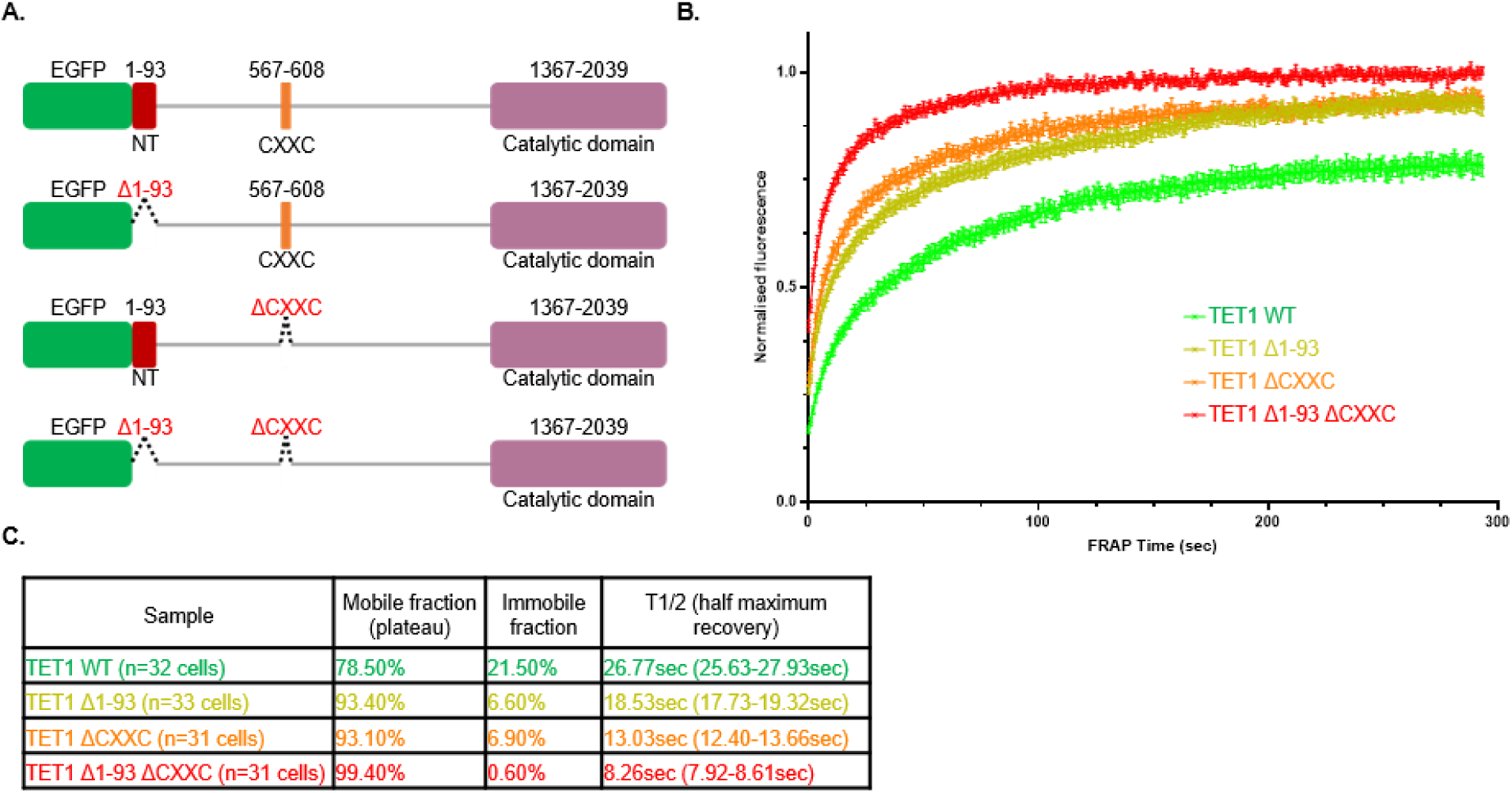
TET1 NT and CXXC act cooperatively to mediate efficient chromatin binding in vivo. **A.** Diagram showing TET1 wild-type and mutant constructs cloned into expression plasmids (EGFP fluorescent reporter) for Fluorescence Recovery After Photobleaching (FRAP) analysis. **B.** Fluorescence recovery of EGFP-TET1 wild-type and mutant constructs following FRAP analysis in mouse embryonic stem cells. Error bars: SEM. **C.** Quantification of FRAP data shown in panel B. For extrapolated half maximum recovery times, confidence intervals are indicated within brackets. The total number of analysed cells is indicated directly in the table for each sample.

### Cryo-EM structure of the nucleosome bound TET1 N-terminal region

The robust chromatin association exhibited by TET1 NT *in vivo* prompted us to test whether this domain directly engages nucleosomes. The nucleosome constitutes the fundamental repeating unit of chromatin, and other factors displaying similar mitotic chromatin retention bind to nucleosomes directly ^57^. We therefore tested this hypothesis using recombinant TET1 NT region and reconstituted nucleosome core particles (NCPs) *in vitro*. To define the molecular basis of this interaction, we determined the cryo-EM structure of TET1 (residues 1-93) bound to the NCP at an overall resolution of 2.47 Å (Fig.4A, S3, S4). We could confidently build a model into the cryo-EM map and assign regions of TET1 to the extra-nucleosomal density. The reconstruction reveals that residues 71-77 of TET1 form a distinct loop that projects into a negatively charged canyon formed between histones H2A and H2B termed the acidic patch (Fig.4B, 4C). Beyond this region, TET1 extends along the C-terminal α-helix of histone H2B (Fig.4A). A second TET1 molecule is observed on the opposite face of the nucleosome, engaging the symmetry-related acidic patch through an equivalent binding mode.

**Figure 4.**
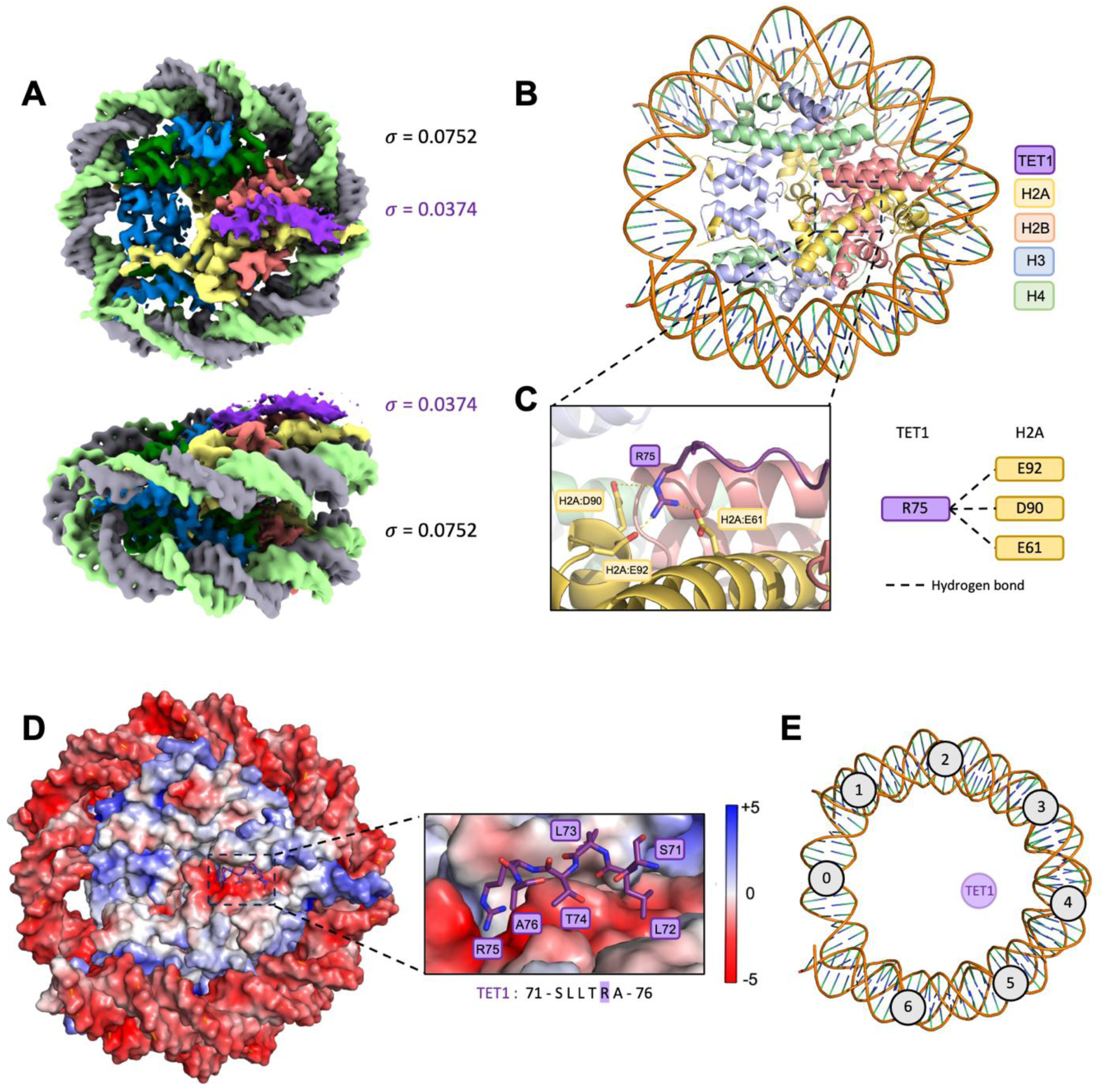
Cryo-EM structure of TET1 in a nucleosome bound state. **A.** Top and side views of the final refined TET1-NCP 3D reconstruction. Nucleosome densities are coloured as follow: H2A (yellow), H2B: (red), H3 (blue), H4 (dark green), TET1 (purple) and DNA (grey/green). Two map thresholds are indicated to distinguish TET1 from histone densities (TET1 σ is purple and NCP σ is black). **B.** Ribbon representation of TET1 bound to the nucleosomal acidic patch. TET1 is shown in purple, DNA in orange and histones H2A, H2B, H3, and H4 in yellow, red, light blue, and green, respectively. **C.** Close-up views of the interactions between TET1 R75 and H2A (left panel). TET1 residues of interest are shown in stick representation. The right panel illustrates the contact points between TET1 R75 and histone H2A. **D.** Electrostatic surface representation of the TET1-NCP complex (blue, +5 kT; white, neutral; red, −5 kT). The right panel shows a zoomed view of the acidic patch: amino acids 71-75 of TET1 are shown in stick representation, with carbon atoms in cyan, oxygen in red, nitrogen in blue. **E.** Position of TET1 on the NCP, relative to nucleosomal superhelical locations.

The TET1 71-77 loop engages the acidic patch through a canonical arginine anchor centred on R75 (Fig.4B, 4C). R75 inserts deeply into the arginine-anchor binding pocket, interacting with H2A residues E61, D90, and E92 (Fig.4D). This interaction represents the most prominent and best-resolved feature of the interface, consistent with the high local resolution observed in this region consistent with its high local resolution (Fig.S3B). According to established nomenclature ^58^, R75 occupies the canonical arginine anchor position commonly used by acidic-patch-binding factors. Additional contacts further stabilize the interface through interactions with histones H2A and H2B: L73 interacts with H2B H109 and E113, L72 contacts H2A E56 and H2B Q47, and S71 engages H2B K116 (Fig.4D).

Beyond the arginine anchor, TET1 extends along the H2B C-terminal α-helix and projects toward nucleosomal DNA at superhelical location +4.5 (SHL +4.5; Fig.4A, 4E). Density corresponding to this region is less well resolved, consistent with its lower local resolution and the presence of a lysine-rich region, that precluded detailed atomic modelling. Nevertheless, the density extends toward nucleosomal DNA at SHL +4.5 and approaches the N-terminal tail of histone H2A near residue K15 (Fig.4A). The remainder of the protein is not resolved in the structure. Together, these findings reveal that TET1 directly recognizes the nucleosome through a canonical acidic-patch arginine anchor that is complemented by an extended interaction along the H2B surface ^58^.

### Functional characterization of TET1 arginine anchor motif

To validate the nucleosome-binding mode observed in our cryo-EM structure, we examined the interaction of TET1 with different nucleosome variants (Fig.S5A). Recombinant TET1 NT (residues 1-93) (Fig.S5B) displayed high affinity binding to NCPs as measured by fluorescence anisotropy assay (Fig.5A). Substitution of the arginine anchor residue with an oppositely charged amino acid (R75E) reduced nucleosome binding by approximately tenfold (Kd=2520nM for R75E mutant versus Kd=276nM for wild-type), confirming the central contribution of R75 to the interaction. Consistent with this result, reduction in the charge of the acidic patch of the nucleosome (acidic patch mutant: H2A^E61A/E91A/E92A^ and H2B^E105A^) greatly reduced TET1 interaction in both pull-down (Fig.5B) and electrophoretic mobility shift assays (EMSA, Fig.S5C), again with dependency on Arg-75 of TET1. Together, these findings validate the acidic-patch interaction observed in the cryo-EM structure and establish R75 as the principal arginine-anchor residue mediating nucleosome recognition.

**Figure 5.**
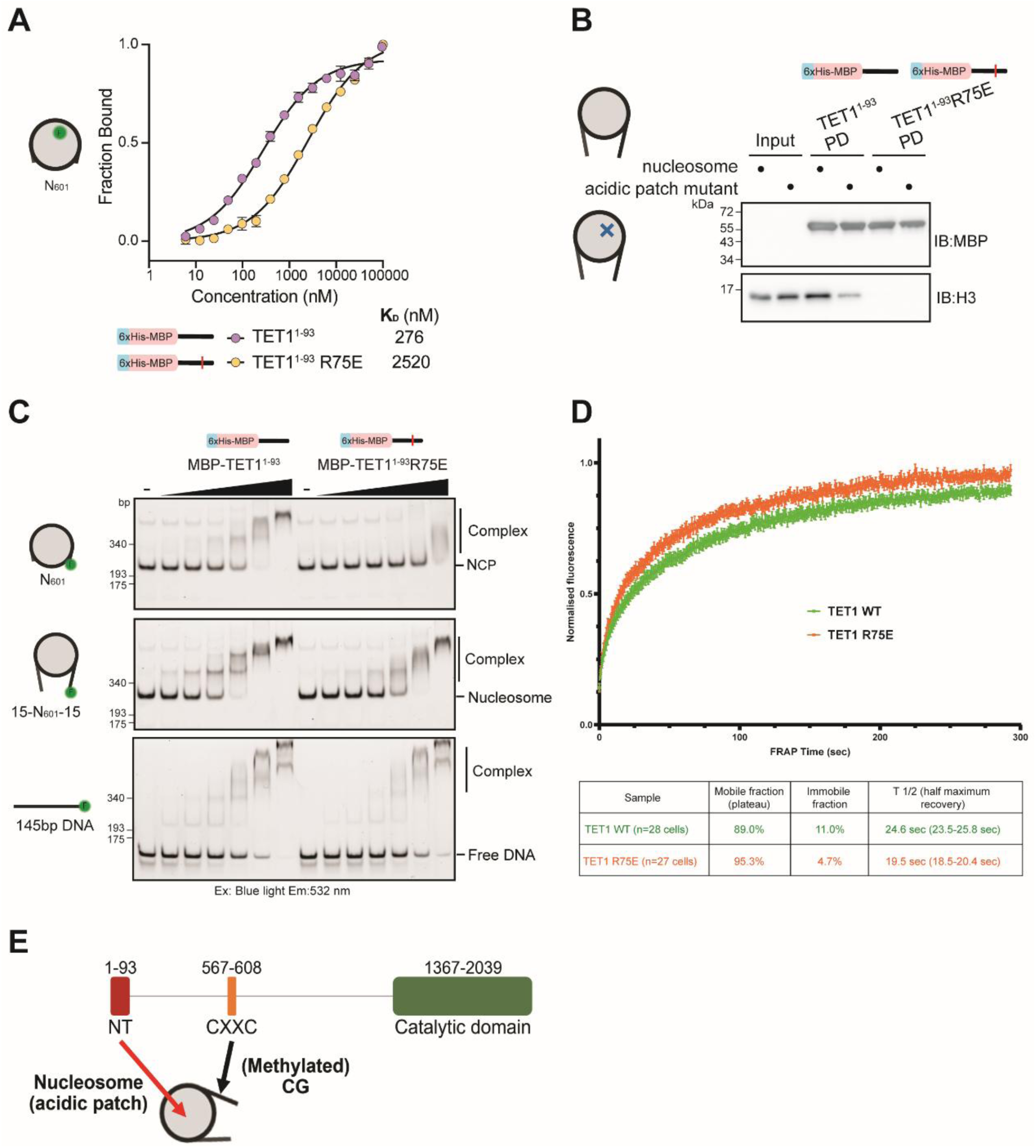
The residue Arg75 mediates a critical interaction with the acidic patch of nucleosomes. **A.** Fluorescence anisotropy assays comparing interaction of TET1 N-terminal region (1-93) wild-type and R75E mutant with fluorescently labelled recombinant NCPs. Error bars represent the standard error of mean (SEM). **B**. Pull down assay to investigate the effects of mutations in both TET1 arginine anchor (R75E) and the acidic patch region of the nucleosome (H2A^E61A/E91A/E92A^ and H2B^E105A^)**. C.** EMSAs comparing binding of TET1 N-terminal region (1-93) wild-type and R75E mutant to different constructs. 5’ FAM (F) labelled DNA was used to monitor NCPs (N601), nucleosomes (15-N601-15) or DNA interaction. Limiting amounts (2.3 nM) of fluorescent labelled tracer were incubated with increasing concentrations (0-1000 nM) of TET1. Complexes were resolved by native-PAGE and imaged using blue light excitation and 532nm emission filters. **D.** Fluorescence recovery of EGFP-TET1 wild-type and mutant (R75E) constructs following FRAP analysis in mouse embryonic stem cells. Error bars: SEM. Quantification of FRAP data is shown below the graph. For extrapolated half maximum recovery times, confidence intervals are indicated within brackets. The total number of analysed cells is indicated directly in the table for each sample. **E.** New working model for TET1 chromatin binding activity, showing the cooperative action of TET1 NT and CXXC domain within the extensive N-terminal regulatory region. The NT mediates multivalent interactions with the nucleosome acidic patch and DNA, while the CXXC recognises cytosines in the context of both methylated and unmethylated DNA.

Interestingly, residual binding remained detectable for both the TET1(R75E) variant and the acidic patch mutant nucleosome (Fig.5A, 5B, S5C), indicating that additional contacts contribute to the interaction, including perhaps nucleosomal DNA (Fig.4A). Consistent with this possibility, TET1 NT bound more efficiently to nucleosomes containing linker DNA than to linker –less NCPs (Fig.5C). Moreover, the NT region displayed appreciable binding to free DNA in EMSA assays, and this activity was only modestly affected by the R75E substitution (Fig.5C). These observations suggest that, in addition to acidic-patch recognition through R75, other residues within the NT region contribute to nucleosome association through multivalent DNA interactions. The DNA binding is likely driven by electrostatic-mediated interactions and as such dispersed across the multiple exposed DNA binding sites in the nucleosome core and linker DNA, and thus not resolved in our cryo-EM structure (Fig.4A).

To assess the impact of the arginine anchor in the context of the full-length TET1 protein *in vivo*, we transfected ESCs with a plasmid encoding for EGFP-tagged TET1 with the R75E point mutation, and subsequently performed FRAP analysis (Fig.5D). This revealed a significantly decreased affinity for chromatin, characterised by shorter recovery times compared to wild-type TET1 (t1/2 ∼19.5sec for TET1(R75E) versus t1/2 ∼24.6sec for the wild-type control). However, the defect caused by this point mutation appeared to be less pronounced than that observed following deletion of the entire NT region (TET1 Δ1-93, Fig.3B, 3C). These findings are consistent with biochemical and structural data, showing additional contributions of other residues within the NT region to chromatin binding. Overall, our study has uncovered the TET1 NT as a multivalent chromatin-binding module that combines acidic-patch recognition with additional DNA-dependent interactions (Fig.5E).

## Discussion

In this study, we uncovered the N-terminal region (NT) as a previously unrecognized functional module within the epigenetic regulator TET1 that mediates chromatin association. Our findings are consistent with earlier reports showing that exogenously expressed TET1 N-terminal fragments lacking the CXXC domain retain chromatin-binding activity and can induce large-scale alterations in nuclear organization when overexpressed ^33,59^. In parallel, a recent proteomic analysis of embryonic stem cell extracts identified TET1 among proteins enriched on nucleosomes and markedly depleted upon extensive mutation of the nucleosomal acidic patch ^57^. Despite these observations, the molecular basis of CXXC-independent chromatin binding had remained unresolved, and it was unclear whether the residual chromatin association reflected direct engagement of nucleosomes or indirect recruitment through interacting partners. Here, we mapped a new chromatin-binding region within the first 93 amino acids of TET1 and demonstrate a direct interaction with nucleosomes through complementary biochemical, structural, and live-cell imaging approaches. Together, these findings establish the NT as a bona fide nucleosome-binding module and reveal an unexpected mechanism underlying TET1 chromatin engagement.

Notably, this region is unique to TET1 and absent from both TET2 and TET3, suggesting that it may confer biological properties specific to this paralogue. Direct nucleosome engagement through the NT could facilitate chromatin exploration independently of sequence-specific recruitment mechanisms and may contribute to previously reported functions of TET1 in cellular reprogramming and pluripotency maintenance ^60–63^. Intriguingly, the NT appears evolutionarily restricted to mammalian species. This observation is noteworthy in light of the divergent biological requirements for TET1 across vertebrates. TET1-deficient zebrafish develop to adulthood without overt developmental defects ^64,65^, whereas TET1 is entirely absent from amphibian genomes such as *Xenopus* ^30^. The acquisition of an NT-mediated nucleosome-binding activity in mammals therefore coincides with the emergence of prominent TET1 functions during peri-implantation development and germline specification ^19,20,23–26,60^. Although causality remains to be established, the evolutionary appearance of this domain is consistent with tight epigenetic control during early mammalian development.

A striking finding of our study is that the NT region promotes TET1 retention on mitotic chromatin. Early models proposed that chromosome condensation leads to widespread eviction of regulatory proteins during mitosis, with only a restricted subset of factors remaining bound to “bookmark” regulatory elements for rapid transcriptional reactivation following cell division ^66–68^. More recent work has challenged this paradigm, revealing that numerous transcription factors and chromatin regulators maintain substantial chromatin association throughout mitosis ^52,55,56^. Although all components of the DNA methylation machinery (DNMT1/3A/3B) remain associated with mitotic chromosomes ^55,69–71^, the behaviour of TET demethylases during this stage of the cell cycle had not been determined. Our results demonstrate that TET1 remains chromatin-bound throughout mitosis and identify the NT as a major determinant of this localization. One attractive possibility is that mitotic retention enables TET1 to preserve regulatory elements in a hypo-methylated state across cell divisions, thereby contributing to epigenetic memory in pluripotent cells. However, given the rapid dynamics of mitotic transitions, directly testing such a model would require new experimental approaches ^72^. Alternatively, NT-mediated nucleosome recognition may primarily function as a general chromatin-tethering mechanism that enhances TET1 occupancy throughout the cell cycle, with mitotic retention representing a natural consequence of this broader activity.

Our structural analyses reveal that TET1 engages the nucleosomal acidic patch through a conserved arginine anchor centred on R75. To our knowledge, this represents the first direct structural characterization of nucleosome recognition by a member of the TET family. The acidic patch is a major interaction hotspot on the nucleosome surface and remains accessible even within highly condensed mitotic chromatin. Recognition of this interface therefore provides a plausible explanation for the differential behaviour of TET1 and TET2 during mitosis. Similar acidic-patch-mediated retention mechanisms have been described for proteins such as RCC1 and CENP-C ^73,74^, which, like TET1, use a surface-exposed arginine residue to engage the canonical acidic-patch pocket. Interestingly, components of the DNA methylation machinery, such as DNMT3 enzymes also exploiting this nucleosomal surface through arginine-anchor motifs ^75–78^, suggesting that acidic-patch recognition may represent a recurrent strategy used by both DNA methylation and demethylation pathways to maintain access to chromatin.

Importantly, the interaction of TET1 extends beyond the canonical acidic-patch pocket. Cryo-EM density reveals additional contacts along the C-terminal α-helix of H2B and toward nucleosomal DNA, indicating a more extensive mode of nucleosome recognition than that observed for compact acidic-patch binders such as LANA ^79^. Similar utilisation of the H2B C-terminal surface has been reported for multiple chromatin-associated complexes, including DNA methylation factors and other nucleosome-binding proteins ^58,75,78,80–84^. These observations suggest that TET1 combines canonical acidic-patch recognition with auxiliary interactions involving broadly accessible nucleosomal surfaces, thereby increasing binding avidity while preserving compatibility with diverse chromatin environments.

Our data support a model in which TET1 integrates multiple modes of chromatin recognition, combining direct nucleosome engagement through the NT with supplementary DNA binding mediated by the NT and CXXC domain ^31–33^ and additional low-affinity interactions with chromatin-associated factors ^45–49^. This multivalent architecture may reconcile the intrinsically weak affinity of TET1 for methylated cytosines with its robust occupancy at regulatory elements observed *in vivo* ^34–39,49^. Consistent with this model, TET1 displays unusually stable chromatin association, evidenced by its nuclear pattern “resistant” to chemical fixation and slow FRAP recovery kinetics relative to most DNA-binding proteins. Our mutational analyses further demonstrate that the NT and CXXC domains act cooperatively to support efficient chromatin binding, highlighting nucleosome engagement as an integral component of TET1 recruitment rather than a secondary accessory interaction.

More broadly, our findings establish direct nucleosome recognition as a defining molecular feature of TET1. By anchoring to the nucleosomal acidic patch, TET1 gains privileged access to chromatin independently of sequence-specific recruitment mechanisms, providing a physical framework for its role in maintaining hypomethylated genomic landscapes in pluripotent cells. These results shed light on the mechanisms by which DNA (de-)methylation defines cellular identity during development.

## Materials and Methods

### Cell culture

All cell lines described in this study were derived from E14Tg2a mouse embryonic stem cells (wild-type parental line) ^85^. *Tet1^Flag^* and *Tet2^Flag^* cell lines are described in a previous publication ^22^, while *Tet1^EGFP^* and H2B-TdT expressing cells were generated in this study.

Cell culture was performed as previously described ^15,22,49^. Cells were incubated in a 37°C, 5% CO_2_ incubator and routinely cultured in serum/LIF medium on gelatin coated plates. Composition of the serum/LIF medium: Glasgow minimum essential medium (GMEM; Sigma, cat. G5154), 10% fetal bovine serum, 1× L-glutamine (Invitrogen, cat. 25030-024), 1x pyruvate solution (Invitrogen, cat. 11360-039), 1× MEM non-essential amino acids (Invitrogen, cat. 11140-035), 0.1mM 2-Mercaptoethanol (Gibco ref. 31350010), 100 U/ml LIF (made in-house). For imaging, cells were plated and cultured directly on polymer coverslips (iBidi cat. 81156 or 80286) with the appropriate coating and culture conditions.

### Genetically engineered cell lines

The *Tet1^EGFP^* knock-in cell line was generated by CRISPR/Cas9-mediated homologous recombination using an established protocol ^22,86^. A single guide RNA (sgRNA) targeting *Tet1* STOP codon was designed using an online tool (http://crispr.mit.edu/) and cloned into Cas9/sgRNA co-expression plasmids (pX459; Addgene cat. 62988). A repair template for homologous recombination carrying EGFP (preceded by a short flexible linker) in frame with TET1 coding sequence was cloned into a pBluescript plasmid backbone by Gibson assembly ^87^. To prevent unwanted cutting of targeted alleles by CRISPR/Cas9, the repair template was designed so that the sgRNA site is disrupted after homologous recombination. Around 1 × 10^6^ E14Tg2a ESCs were transfected with both the Cas9/sgRNA plasmid and the repair template using Lipofectamine 3000 (ThermoFisher Scientific, cat. L3000008) and following the manufacturer’s instructions. After 48 h, single EGFP-positive cells were sorted into wells of 96-well plates by flow cytometry. ESC clones were further expanded in 24-well plates and genomic DNA was extracted (Qiagen, cat. no. 69506) for genotyping using primer pairs specific to the wild-type or the EGFP-tagged allele. One of the primers was located outside of the homology arms of the repair template to ensure proper homologous recombination within endogenous *Tet1* locus. The desired modification was confirmed by Sanger sequencing of PCR products for both alleles. Homozygous *Tet1^EGFP^* ESC clones were further expanded and stored in liquid nitrogen for long-term storage.

To generate H2B-TdT expressing cells, around 1 × 10^6^ E14Tg2a ESCs were transfected using Lipofectamine 3000 (ThermoFisher Scientific, cat. L3000008) with a mammalian expression plasmid encoding for Histone H2B fused to a tdTomato fluorescent reporter (H2B-TdT) under a constitutive promoter, also including a puromycin resistance cassette. Following 48h, transfected cells were selected with 1µg/ml puromycin for at least one week to enrich the population in cells carrying a random integration of the transgene. Finally, a bulk population of tdTomato-positive cells was purified by flow cytometry. H2B-TdT expressing cells were further expanded and stored in liquid nitrogen for long-term storage.

### Immunofluorescence

The protocol used for immunofluorescence was described previously ^22,88^. Cells were washed with PBS and fixed with 4% PFA for 10 min at room temperature. After fixation, cells were washed with PBS and permeabilised with a solution of PBS containing 0.3% (vol/vol) Triton X-100 for 10 min at room temperature. Samples were blocked in blocking buffer (PBS supplemented with 0.1% (vol/vol) Triton X-100, 1% (wt/vol) BSA, and 3% (vol/vol) goat serum) for at least 1 h at room temperature. After blocking, samples were incubated with primary antibodies diluted at the appropriate concentration (see Antibodies table) in blocking buffer overnight at 4°C. After four washes with PBS containing 0.1% (vol/vol) Triton X-100, samples were incubated with fluorescently labelled secondary antibodies (ThermoFisher Scientific, cat. A32728TR or cat. A32731TR) diluted 1:500 in blocking buffer for 1 h at room temperature in the dark. Cells were washed four times with PBS containing 0.1% (vol/vol) Triton X-100. DNA was stained with DAPI for 5 min at room temperature, and cells were submitted to a final wash with PBS for 5 min. Immunofluorescence samples were imaged using either a brightfield (Nikon Ti) or a spinning disk (Nikon CSU-X1) microscope. Images were processed and analysed using the software Fiji.

Immunofluorescence data was used for the quantification of TET1/2 co-localisation with mitotic chromatin. In all fields of view, mitotic cells were counted and identified visually due to the typical DNA condensation observed in pro-metaphase, metaphase or anaphase. The DNA (DAPI) and TET1/2 (Flag) channels were investigated concomitantly in order to assess the overlap between the two fluorescence signals. Mitotic cells were classified into four categories: no TET expression, no co-localisation, partial co-localisation or full co-localisation of TET1/2 with chromatin. This analysis was not blinded and samples were not randomised. Data was plotted using the software GraphPad Prism.

### Live-cell imaging and FRAP analysis

Live-cell imaging and FRAP analysis were performed as described previously ^89^. With the exception of the *Tet1^EGFP^*knock-in cell line (endogenous fluorescence), cells were cells were transfected with expression plasmids encoding for EGFP-tagged TET1:2 constructs of interest using Lipofectamine 3000 (ThermoFisher Scientific, cat. L3000008) and following the manufacturer’s protocol. Cells were analysed the next day following a medium change. For live-cell imaging, cells were imaged using either a brightfield (Nikon Ti) or a spinning disk (Nikon CSU-X1) microscope at 37 °C with 5% CO_2_. Images were processed and analysed using the software Fiji.

FRAP was performed using the Zeiss LSM 880 confocal microscope at 37 °C with 5% CO_2_. For each cell, EGFP-TET1 signal was imaged every 1 s for 300 s with five images recorded before bleaching a selected spot (FRAP spot) with 100% laser power. Images were acquired using the Zeiss ZEN software (black edition) and processed using a custom script with Fiji software (https://doi.org/10.5281/zenodo.2654601) ^89^. Fluorescence was measured at the bleached spot (FRAP spot), as well as a non-bleached control region (control spot) to account for photobleaching during the experiment. Additionally, the fluorescence outside of transfected cells was measured as background. The first time point (*T*_0_) was defined as the first post-bleach image. For each time point, the FRAP fluorescence signal was normalised to fluorescence values before photobleaching. A ’Two phase association’ model (nonlinear regression) was used to fit experimental data using the software GraphPad Prism. The plateau (fluorescence at the last time point of the FRAP experiment), corresponding to the mobile fraction, was interpolated from the fitted curve. Conversely, the immobile fraction corresponds to the pool of fluorescent proteins not recovered during the experiment (=100% − mobile fraction). The half maximum recovery times (t1/2) was also interpolated from the fitted curve. Data was plotted using the software GraphPad Prism.

### Protein expression and purification

The cDNA encoding mouse TET1(1–93) was subcloned into the pLIC vector (Addgene # 29656) in frame with a N-terminal TEV-cleavable 6xHis-MBP tag. For structural studies TET1(1-93) was expressed in *E. coli* Lemo21(DE3) cells, grown in Terrific Broth at 37 °C to OD₆₀₀ = 0.6, then induced with 0.2 mM IPTG for 16 h at 19 °C. Cells were harvested, resuspended in lysis buffer [50 mM Tris pH 8.0, 500 mM NaCl, 0.1% (v/v) NP-40, 5 mM 2-mercaptoethanol], lysed by sonication, and clarified by centrifugation. The supernatant was incubated with Nickel-sepharose resin (Cytiva) for 1 h at 4 °C, washed with lysis buffer containing 1 M NaCl, then equilibrated in 300 mM NaCl. 6xHis-MBP-TET1(1–93) was eluted with 400 mM imidazole, dialyzed, and TEV-cleaved on beads for 16 h at 4 °C. The untagged protein was purified by ion-exchange chromatography (Resource S, GE Healthcare), and peak fractions were concentrated to about 4 mg/mL.

For biochemical assays, 6xHis-MBP-TET1(1-93) and the R75E mutant were expressed in BL21 (DE3 RIL) *E. coli* cells for 3 hours with 1 mM IPTG at 30°C in 2xYT broth. Cell pellets were resuspended in lysis buffer (50 mM HEPES pH 7.5, 300 mM NaCl, 10% (v/v) glycerol, 2 mM β-mercaptoethanol, 1X protease inhibitor cocktail (2.2 mM PMSF, 2 mM benzamidine HCl, 2 μM leupeptin, 1 μg.mL^−1^ pepstainA), 4 mM MgCl_2_, 5 μg.mL^−1^ DNAse and 500 μg.mL^−1^ lysozyme) and nutated at 4°C for 30-60 minutes, before additional lysis using a sonicator (2 s on, 2 s off for total 20 s at 50% amplitude). Bacterial cell debris was pelleted by centrifugation for 25 minutes at 39,000 x g at 4°C and clarified lysate was filtered through a 0.4 μm filter. 6xHis-MBP-TET1^1–93^ clarified lysate was supplemented with 10 mM imidazole before being applied to a HiTrap chelating HP column (Cytiva) pre-charged with nickel ions, extensively washed with wash buffer (50 mM HEPES pH 7.5, 300 mM NaCl, 10% (v/v) glycerol, 10 mM imidazole, 2 mM β-mercaptoethanol, 0.1 mM AEBSF) and eluted with elution buffer (50 mM HEPES pH 7.5, 300 mM NaCl, 10% (v/v) glycerol, 300 mM imidazole, 2 mM β-mercaptoethanol, 0.1 mM AEBSF) over a 15 CV gradient. Fractions containing 6xHis-MBP-TET1^1–93^ were pooled and diluted to 150 mM NaCl in SP buffer No Salt (10 mM HEPES pH 7.0, 1 mM DTT, 5% (v/v) glycerol, 0.1 mM AEBSF), before further purification by ion exchange chromatography. Diluted 6xHis-MBP-TET1^1–93^ fractions were applied to a pre-equilibrated SP HP column and washed extensively with SP A buffer (10 mM HEPES pH 7.0, 100 mM NaCl, 1 mM DTT, 5% (v/v) glycerol and 0.1 mM AEBSF), before eluting with SP B buffer (10 mM HEPES pH 7.0, 1000 mM NaCl, 1 mM DTT, 5% (v/v) glycerol and 0.1 mM AEBSF), across an 18 CV gradient. Fractions containing 6xHis-MBP-TET1^1–93^ were then pooled, concentrated in a 10 kDa MWCO centrifugal filter unit, before being applied to a HiLoad Superdex 75 16/600 (Cytiva) pre-equilibrated with gel filtration buffer (10 mM HEPES pH 7.5, 150 mM NaCl, 5% (v/v) glycerol, 1 M DTT, and 0.1 mM AEBSF). Fractions containing 6xHis-MBP-TET1^1–93^ were then pooled, concentrated, aliquoted and flash frozen in liquid nitrogen, before being stored at –80°C.

### Nucleosomal DNA and large-scale nucleosome reconstitution

For the structure, The 147-bp strong positioning 601 DNA sequence ^90^ was used. To produce sufficient quantity of NCPs for structural studies, a large-scale protocol was employed as indicated in a previous study ^84^. Briefly, 601 DNA was generated in milligram amounts from a concatemer plasmid, excised by restriction digestion, and size-selected ^91^. Native *X. laevis* histones were co-expressed in *E. coli* and purified as described ^92^. NCPs were assembled by overnight dialysis from 2 M to 0.25 M NaCl and heat-shifted at 37 °C for 120 min.

For biochemistry, Unmodified human histones, as well as variants H2B T115C and acidic patch mutants (H2A E61A E91A, H2B E105A), were expressed and purified as previously described ^93–97^. Histone H3 without any cysteine residues (C91S C110A) was used for all biochemical experiments. Briefly, histones were expressed in BL21 (DE3 RIL) cells and resolubilised from inclusion bodies. Histones were further purified by ion exchange chromatography prior to dialysis in 1 mM acetic acid and lyophilisation.

Labelling of H2B T115C was performed essentially as described before ^97^ with Oregon-Green488 maleimide dye (AAT Bioquest) prior to octamer reconstitution and checked for labelling efficiency by 1D intact weight ESI mass spectrometry (SIRCAMs, School of Chemistry, University of Edinburgh). Histone octamers containing the purified histones were reconstituted as preciously described by ^93,95,96^. Histones were rehydrated to ∼5 mg.mL-1 in unfolding buffer (20 mM Tris pH 7.5, 7 M Guanidine-HCl, 10 mM DTT) for 30 minutes at room temperature. Unfolded histones were mixed together in a 1.2:1.2:1:1 molar ratio of H2A:H2B:H3:H4 and diluted in unfolding buffer to a final concentration of 2 mg.mL-1. The histone mixture was then dialysed extensively in refolding buffer (15 mM Tris pH 7.5, 2 M NaCl, 1 mM EDTA, 5 mM β-mercaptoethanol) O/N at 4°C. Refolded octamers were separated from soluble aggregates, tetramers, dimers and unpaired histones by size exclusion chromatography on a Superdex S200 16/60 (Cytiva) in refolding buffer. Peak fractions corresponding to the refolded octamers were pooled, concentrated and then used immediately for nucleosome reconstitution, or stored at –20°C after addition of 50% (v/v) glycerol.

Unlabelled and FAM labelled Widom 601 DNA for nucleosome reconstitution were generated by PCR and purified essentially as previously described ^75,90,94,96–98^. And either consisted of 145bp 601 core 601 fragment (N601) or core fragment with two symmetrical linear linker sequences (15-N601-15). Fluorescent dyes were incorporated in the primers (IDT technologies). Labelled and unlabelled nucleosomes were reconstituted as essentially previously described ^75,94–98^.

### Cryo-EM sample preparation

TET1 (1-93) and nucleosomes used in the preparation of the complex for cryo-EM studies were produced as indicated above. 147-bp nucleosomes and TET1 (aa1-93) were mixed at a molar ratio of 1:3 (NCP:TET1). The complex was incubated at 4°C for 1 h before being purified by size exclusion chromatography (Superose 6 Increase 3.2/300). To prepare the cryo-EM grids, the freshly purified complex was first cross-linked with 0.1% glutaraldehyde, then, immediately, 5 μl of sample were applied on Quantifoil R 1.2/1.3 Gold foil on Gold 300 mesh grid, plasma cleaned on a model 1070 NanoClean (Fischione Instruments) for 90 s at 30% power and with a mixture of 80% argon and 20% oxygen. After application the grids were blotted for 4 s at a force setting of 5 and then plunged into liquid ethane using a Vitrobot mark IV (FEI) system with a chamber humidity of 95% at 10°C.

Images were collected on a 300 kV Titan Krios (FEI) at the Institut de Génétique et de Biologie Moléculaire et Cellulaire (IGBMC) equipped with a Falcon 4i camera and Selectris energy filter in super resolution counting mode using SerialEM software ^99^ with a physical pixel size of 0.729 Å and a target defocus range from −0.8 to −2.3 μm. Exposures of 2.51 s were dose-fractionated into 801 EER frames with a dose rate of 0.064 electrons pixel−1 s−1, with a total dose of 50.25 e−/Å2 (Table 1). A total of 19 285 movies were collected.

**Table 1:** Refinement and model statistics for the reconstruction of TET1 (1-93) with the NCP Data Processing.

| Particles | Extracted | Used for 3D | Final map |
|---|---|---|---|
|  | 3 713 496 | 1 114 075 | 279 302 |
| Pixel size | 0.862 |  |  |
| Defocus range (µm) | -0.8 to -2.3 |  |  |
| Voltage (kV) | 300 |  |  |
| Total Electron dose (e<sup>-</sup>/Å<sup>2</sup>) | 50.25 |  |  |
| Nominal resolution (Å) | 2.47 |  |  |
| cFAR | 0.57 |  |  |
| Directional resolution range (Å) | 2.28 – 2.81 |  |  |
| Map sharpening B-factor (Å<sup>2</sup>) | -31.7 |  |  |
| Modeling |  |  |  |
| Atomic modeling Refinement package | Phenix, Coot, Chimera |  |  |
| PDB ID / EMDB ID | **** / EMD-**** |  |  |
| Model composition | Protein | Ligand | DNA |
|  | 773 | 0 | 290 |
| CCVolume | 0.81 |  |  |
| CCmask | 0.80 |  |  |
| Model-to-map FSC 0.5 (Å) | 2.8 |  |  |
| B-Factors | Protein | Ligand | Nucleotide |
|  | 111.78 | 0 | 182.97 |
| RMSD | Length (Å) (# > 4 σ) | Angles (°) (# > 4 σ) |  |
|  | 0.004 | 0.640 |  |
| MolProbity score | 1.44 |  |  |
| Clash score | 8.15 |  |  |
| Poor Rotamer (%) | 0.78 |  |  |
| Ramachandrans (%) | Favored | Allowed | Outliers |
|  | 98.01 | 1.99 | 0 |
| CaBLAM outliers (%) | 0.27 |  |  |
| EMRinger score | 3.29 |  |  |

### Cryo-EM data processing and model building

The images in the dataset were aligned and corrected for beam-induced motion using MotionCor2^100^. Contrast transfer function (CTF) parameters were estimated using CTFFIND4 ^101^. All subsequent data processing was performed in CryoSPARC ^102^. Micrographs were denoised in cryoSPARC prior to particle picking, and the denoised micrographs were used for the initial particle picking with the Blob Picker. Low-quality and false particles were removed by 2D classification. The best 2D classes were then selected to generate two ab initio models: one derived from high-quality particles and a second generated from discarded particles, using a limited number of iterations. These models were subsequently used as references for heterogeneous refinement to further clean the dataset and retain particles corresponding to the complex of interest.

The selected particles were then used for template-based particle picking, allowing a more specific particle selection and substantially increasing the total number of particles to 3,713,496. The newly extracted particles were subjected to 2D classification, and the best classes were used for a second round of heterogeneous refinement, resulting in 1,114,075 particles. The resulting particle set was subjected to global CTF refinement, followed by non-uniform refinement and 3D classification to assess the conformational heterogeneity. The best particles subset, comprising 279 302 particles, was then selected for homogenous refinement. The final reconstruction reached an overall resolution of 2.47 Å at the FSC = 0.143 criterion.

Initial particle processing and refinements were performed in C1 symmetry. After confirming that the TET1-associated density did not display any obvious asymmetric features, C2 symmetry was applied during the final refinement to improve the overall map resolution.

The initial model of the TET1(1-93)-nucleosome complex was predicted using AlphaFold ^103^. However, the predicted TET1(1-93) structure did not fully match the extra density observed in the cryo-EM map, as AlphaFold predicted a C-terminal α-helix that was not supported by the experimental density. The most confident residue in the prediction was Arg75, which was positioned in contact with the acidic patch and corresponded to the observed extra density. Accordingly, the AlphaFold-predicted model was truncated to residues 71–76, corresponding to the region matching the additional density at the acidic patch. Docking into the EM map was performed using UCSF Chimera ^104^. The model was manually adjusted and rebuilt as necessary in Coot ^105^, followed by rigid-body refinement and global minimization in Phenix ^106,107^ (Table 1). The refined model was subsequently subjected to iterative real-space refinement cycles with secondary structure restraints in Phenix ^106,107^, and model errors were inspected and corrected in Coot ^105^. Molecular graphics and structural analyses were performed using PyMOL (PyMOL Molecular Graphics System, version 2.5, Schrödinger, LLC) and UCSF Chimera ^104^.

### Fluorescence anisotropy

Fluorescence anisotropy experiments were performed by incubating 6.1 nM to 100 μM 6xHis-MBP-TET1^1–93^ and 6xHis-MBP-TET1^1–93(R75E)^ proteins with 25 nM Oregon Green-488 labelled NCPs (145bp widom 601, H2A, H2B T115C-OG488, H3.1 C96S C110A, H4). Samples were incubated in FP buffer (15 mM HEPES pH 7.5, 150 mM NaCl, 0.05 mg.mL-1 BSA, 10% (v/v) glycerol, 0.05% (v/v) Triton, 1 mM DTT) for 30 minutes at room temperature. Fluorescence anisotropy at 485/535 nm was detected using a SpectraMax iD5 (Molecular Devices) plate reader. Fluorescence anisotropy data was correct for background fluorescence with buffer only, and anisotropy values (r) were calculated using the formula below, where G = the G factor for the plate reader.

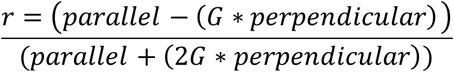

Anisotropy values were converted to fraction bound, using the formula below, where r = the anisotropy value for a given concentration, Af anisotropy value for the nucleosome only sample, and Ab for the anisotropy value for the highest concentration of 6xHis-MBP-TET1^1–93^/R75E mutant protein and nucleosome mixture.

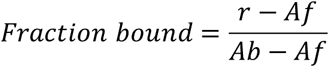

Fraction bound was plotted against TET1 concentration in GraphPad Prism, and fit with a Specific binding with Hill slope curve.

### Nucleosome pulldown

10 μg of 6xHis-MBP-TET1^1–93^ and 10 μg of 6xHis-MBP-TET1^1–93(R75E)^ of was immobilised on 25 μL Amylose Resin (New England Biolabs) in 100 μL pulldown buffer (50 mM HEPES pH 7.5, 150 mM NaCl, 0.02% (v/v) NP40, 0.1 mg.mL^−1^ BSA, 10% (v/v) glycerol, 1 mM EDTA, 1 mM DTT) for 2 h at 4°C with rotation. Beads were washed by centrifugation at 500 x g for 1 minute at 4°C and resuspension in 1 mL of pulldown buffer. Beads were then incubated with 1.25 μg nucleosomes (175bp Widom 601 DNA wrapped) in the same buffer for 2 h at 4°C with rotation. Pulldowns were washed three times as above, using 750 μL of pulldown buffer, before the beads were then resuspended in 2xSDS loading dye and boiled for 5 minutes to release bound proteins. 5% of input nucleosomes were loaded as a control alongside 5 μL of the pulldown reactions. Samples were analysed by western blotting for MBP and H3.

### Electrophoretic mobility shift assays (EMSA)

EMSAs were performed as previously described ^96^. 6-Carboxyfluorescein (5’ 6-FAM) labelled nucleosomes (at 1 ng.uL-1 each) were incubated with various concentration ranges, as noted in figure legends of particular experiment, of 6xHis-MBP-TET1^1–93^ and 10 μg of 6xHis-MBP-TET1^1–93(R75E)^ proteins in EMSA buffer (15 mM HEPES pH 7.5, 75 mM NaCl, 0.05 mg.mL^−1^ BSA, 10% (v/v) glycerol, 0.05% Triton (v/v), 1 mM DTT, 0.5 mg.mL^−1^ competitor DNA (low molecular weight from salmon sperm (Sigma-Aldrich), 8% (w/v) sucrose, 0.01% (w/v) bromophenol blue) to a final volume of 12 μL. Samples were incubated on ice for 1 h, and products were separated by native-PAGE using 1xTris Glycine as running buffer for 90 minutes at 4°C. Gels were imaged for FAM signal (Excitation Blue light, Emission 532nm) using BioRAD ChemiDoc MP.

### Antibodies

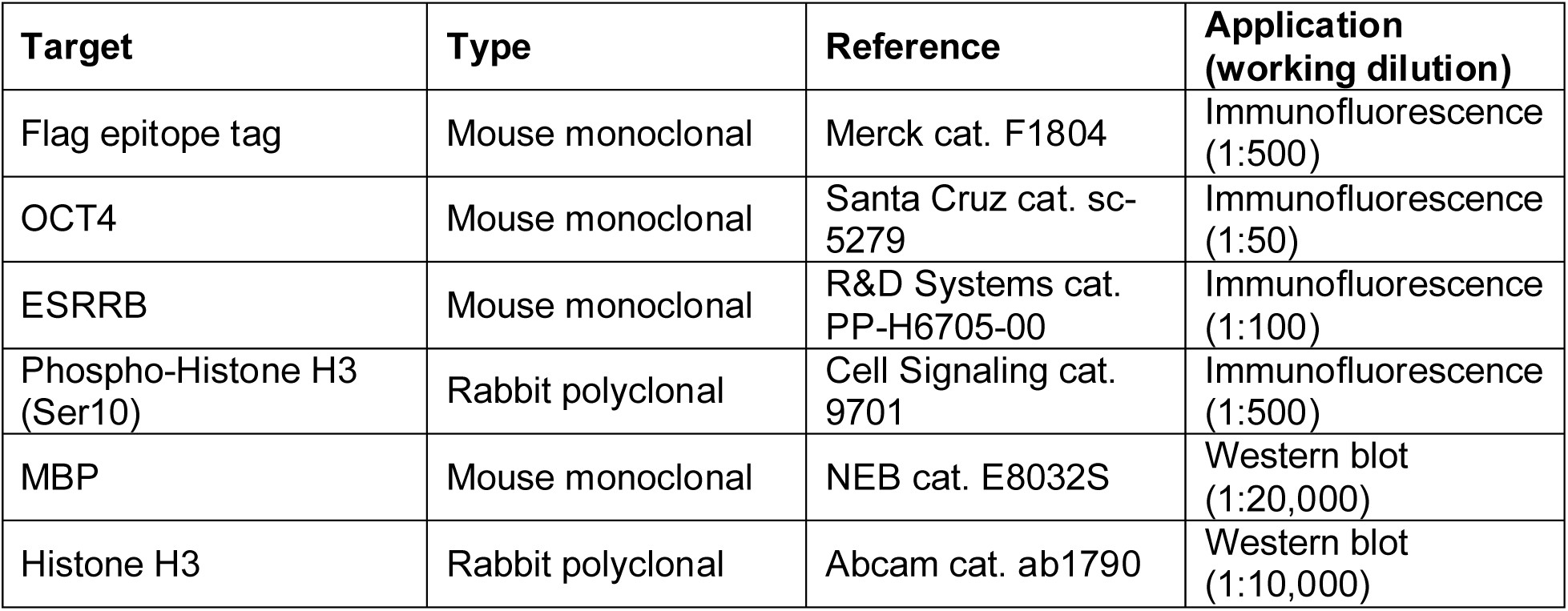

### Oligonucleotides

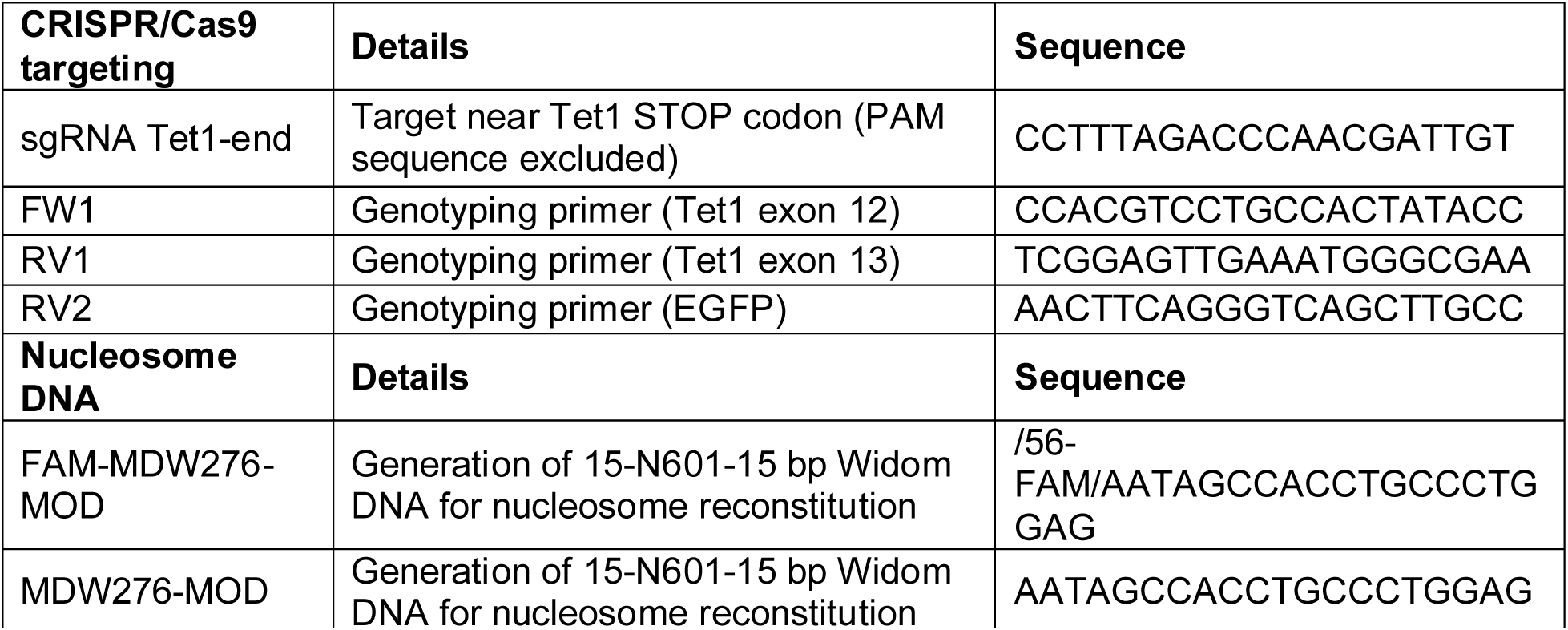
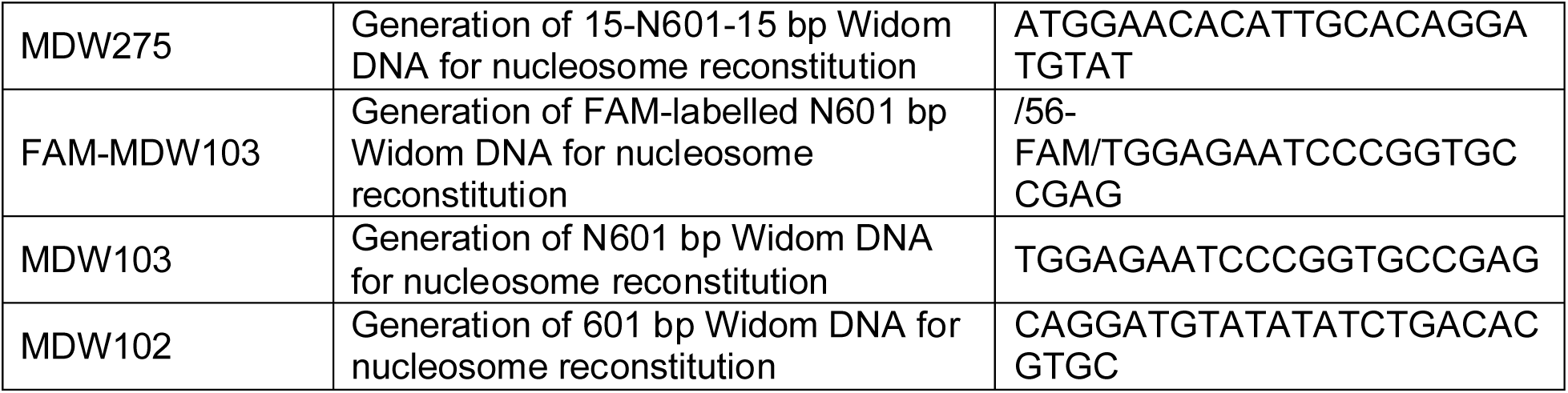

### Plasmids

| **Plasmids** | **Details** |
|---|---|
| sgRNA Tet1 end-Cas9-T2A-Puro | Mammalian expression plasmid encoding for both Cas9 (Puromycin selection) and a sgRNA targeting Tet1 locus |
| Targeting vector_Tet1-EGFP | Targeting vector for CRISPR/Cas9-mediated homologous recombination (Tet1<sup>EGFP</sup> cell line) |
| pPYCAG-EGFP-TET1 WT | Mammalian expression plasmid encoding for TET1 full-length fused with an EGFP fluorescent reporter |
| pPYCAG-EGFP-TET2 WT | Mammalian expression plasmid encoding for TET2 full-length fused with an EGFP fluorescent reporter |
| pPYCAG-EGFP-TET1 delta1-93 | Mammalian expression plasmid encoding for TET1 with a deletion of the first 93 amino acids fused with an EGFP fluorescent reporter |
| pPYCAG-EGFP-TET1 deltaCXXC | Mammalian expression plasmid encoding for TET1 with a deletion of the CXXC domain fused with an EGFP fluorescent reporter |
| pPYCAG-EGFP-TET1 delta1-93 delta CXXC | Mammalian expression plasmid encoding for TET1 with a deletion of both the first 93 amino acids and the CXXC domain fused with an EGFP fluorescent reporter |
| pPYCAG-EGFP-TET1 (R75E) | Mammalian expression plasmid encoding for TET1 full-length with a point mutation (R75E) fused with an EGFP fluorescent reporter |
| pPYCAG-EGFP-TET1 1-856 | Mammalian expression plasmid encoding for truncated TET1 fragment fused with an EGFP fluorescent reporter |
| pPYCAG-EGFP-TET1 1034-2039 | Mammalian expression plasmid encoding for truncated TET1 fragment fused with an EGFP fluorescent reporter |
| pPYCAG-EGFP-TET1 1-108 | Mammalian expression plasmid encoding for truncated TET1 fragment fused with an EGFP fluorescent reporter |
| pPYCAG-EGFP-TET1 1-93 | Mammalian expression plasmid encoding for truncated TET1 fragment fused with an EGFP fluorescent reporter |
| pPYCAG-EGFP-TET1 1-64 | Mammalian expression plasmid encoding for truncated TET1 fragment fused with an EGFP fluorescent reporter |
| pPYCAG-EGFP-TET1 52-108 | Mammalian expression plasmid encoding for truncated TET1 fragment fused with an EGFP fluorescent reporter |
| pLIC-His-MBP-TET1 1-93 | Bacterial expression plasmid encoding for truncated TET1 fragment fused with an 6xHis-MBP affinity tag |
| pLIC-His-MBP-TET1 1-93 (R75E) | Bacterial expression plasmid encoding for truncated TET1 fragment with a point mutation (R75E) fused with an 6xHis-MBP affinity tag |

### Protein alignments and evolutionary conservation analysis

All protein sequences were obtained from the UniProt database ^108^. For comparison of TET1 in mammals, the following protein sequences were used (UniProt protein IDs): A0A6P5J946, F6V6A0, Q3URK3, G3SXN7, Q8NFU7, H2Q1Z7, F1SUI3, A0A8B8VKU6, F1MGK2, A0A452E1W7, A0A671EG23, A0A9L0SMP6, A0ABI7XXE9, A0A8I3MT69, D2GXI8. For comparison of TET1 in amniotes (mammals, reptiles, birds), the following protein sequences were used (UniProt protein IDs): A0A7M4EF40, A0A8C3I386, A0A8V0ZYW0, A0A7L0BLB2, A0A493T2Y8, A0A6J1UE43, H9GVA7, A0A6P5J946, F6V6A0, Q3URK3, Q8NFU7, G3SXN7, A0A671EG23, A0A8B8VKU6. For comparison of TET1 in vertebrates (mammals, reptiles, birds, fishes), the following protein sequences were used (UniProt protein IDs): A0A8M9Q186, A0A6P3WB57, A0A1S3T223, A0A673AZS8, A0A6P8TNS4, A0A3Q1C314, Q3URK3, Q8NFU7, G3SXN7, F1SUI3, A0A671EG23, A0A6P5J946, A0A7M4EF40, A0A8C3I386, A0A8V0ZYW0, A0A7L0BLB2, A0A493T2Y8, A0A6J1UE43, H9GVA7. For comparison of TET1-2-3 paralogues in mouse and human, the following protein sequences were used (UniProt protein IDs): Q8NFU7, Q3URK3, Q8BG87, O43151, Q6N021, Q4JK59. When possible, the longest isoform (containing a CXXC domain) was selected for analysis. Of note, the TET1 paralogue is absent in amphibians.

Protein alignments were performed using the Clustal Omega software ^109^ (https://www.uniprot.org/align) with default parameters. Protein alignment files were further processed and visualised using the software Jalview version 2.11.5.2 ^110^. For analysis of protein conservation, all insertions/gaps in the alignment were ignored to allow direct comparison with the reference sequence (mouse TET1, UniProt protein ID: Q3URK3). Conservation scores at each position in the protein alignment (calculated using the AMAS method with scores range 0-11 ^111^) were exported and visualised as heatmaps using the software GraphPad Prism.

### Reagents and data availability

Request for resources and reagents should be directed to the corresponding authors: Raphaël Pantier, Elisa Bergamin or Marcus Wilson.

Raw and processed data used to generate the Figures will be deposited on a publicly available database at the time of publication.

Cryo-EM coordinates of the TET1-NCP complex have been deposited in the Protein Data Bank (PDB) with accession code 28KG, and the map was deposited in the Electron Microscopy Data Bank (EMDB) with accession code EMD-56575.

## Acknowledgments

We thank Ian Chambers and Adrian Bird for constructive feedback throughout the project. We thank Bertrand Vernay and David Kelly for microscopy support, Claire Cryer and Fiona Rossi for assistance with flow cytometry. The authors wish to thank the members of the Integrated Structural Biology platform at IGBMC for technical assistance.

Work in the Wilson lab is supported by a Sir Henry Dale Fellowship from the Wellcome Trust [210493/Z/18/Z] and Medical Research Council (T029471/1). This work was also supported by the Edinburgh Protein Production Facility (EPPF) and the Centre Optical Instrumentation Laboratory (COIL), which received funding from a core grant (203149) to the Wellcome Centre for Cell Biology at the University of Edinburgh. Raphaël Pantier received funding from the French National Research Agency (ANR-25-CE12-7883 and ANR-25-ERCS-0008) and the University of Strasbourg (IDEX 2025). The work in the Bergamin lab was funded in part by the Fondation pour la Recherche Médicale (grant number AJE20171039021), an ATIP-Avenir award (contract # 188435), an award from La Ligue Contre le Cancer (JMG/SP – n° 01B.2021), startup funds from the IGBMC, ANR-23-CE11-0002-03, ANR-24-CE12-1416, ANR-the French Infrastructure for Integrated Structural Biology (FRISBI) ANR-10-INBS-0005, EquipEx + France cryo-EM (ANR-21-ESRE-0046), salary support by the Centre National de la Recherche Scientifique to E.B and R.P; F.M was supported by a 4^th^ year fellowship from Fondation ARC.

## Authors contribution

**Franck Martin:** Investigation, Methodology, Formal analysis, Visualization, Writing – original draft, Writing – review & editing. **Hayden Burdett:** Conceptualization, Investigation, Methodology, Formal analysis, Visualization, Writing – original draft, Writing – review & editing. **Mélody Matelot:** Investigation, Validation, Writing – review & editing. **Chloé Ngon Baroung:** Investigation, Validation, Writing – review & editing. **Beatrice Alexander-Howden:** Investigation, Writing – review & editing. **Marcus D. Wilson:** Conceptualization, Methodology, Supervision, Funding acquisition, Writing – original draft, Writing – review & editing. **Elisa Bergamin:** Conceptualization, Methodology, Supervision, Funding acquisition, Writing – original draft, Writing – review & editing. **Raphaël Pantier:** Conceptualization, Investigation, Methodology, Formal analysis, Visualization, Supervision, Funding acquisition, Writing – original draft, Writing – review & editing.

## Supplementary material

### Supplementary Figures

**Supplementary Figure 1 (related to Figure 1).**
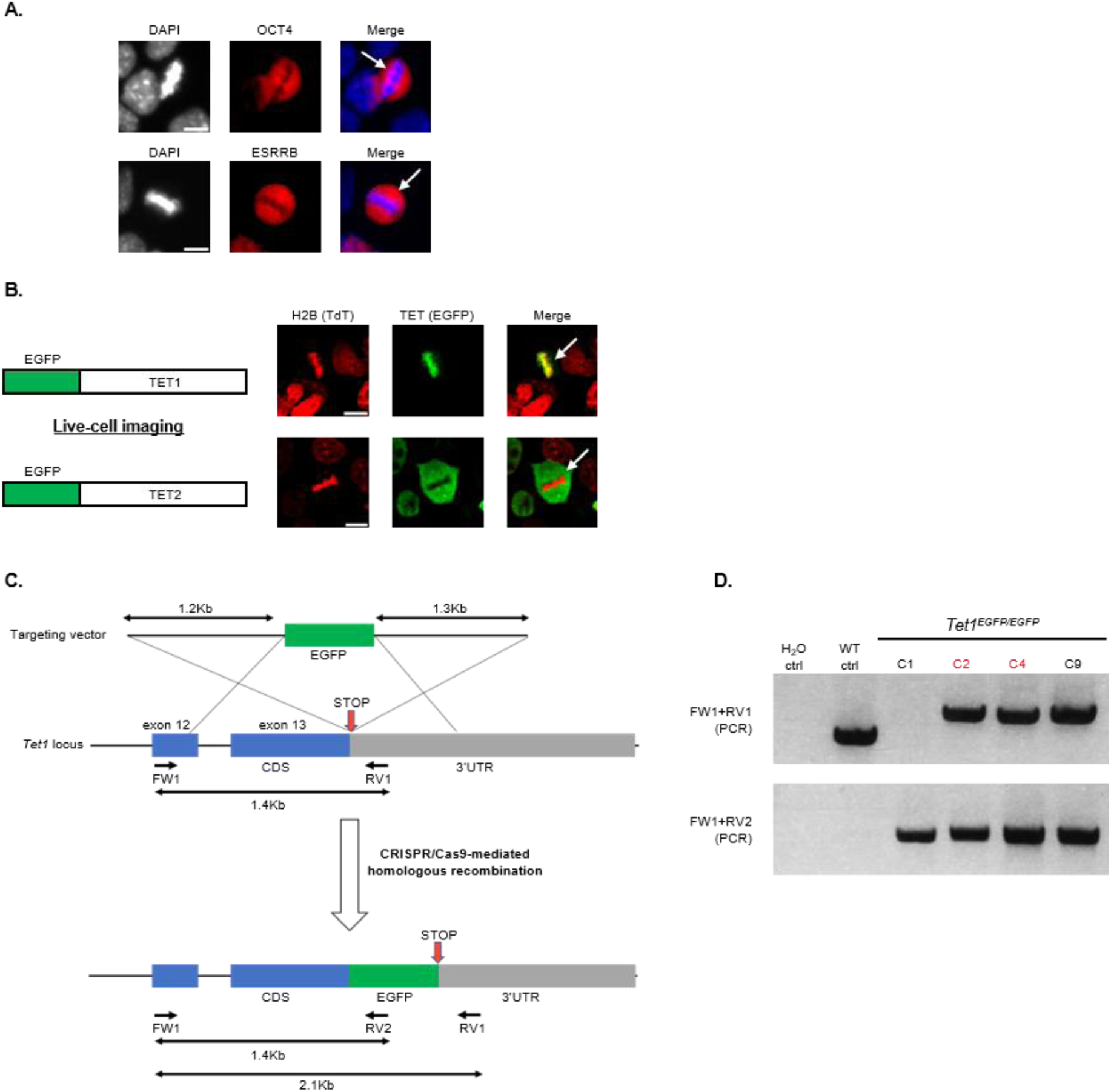
A. Immunofluorescence of endogenous transcription factors OCT4 and ESRRB in mouse embryonic stem cells. DNA was stained with DAPI. Arrows indicate artefactual mis-localisation of these factors in mitotic cells. Scale bars: 10µm. **B.** Live-cell imaging showing the sub-nuclear localisation of wild-type EGFP-TET1 and EGFP TET2 constructs transfected into mouse embryonic stem cells. Chromatin was visualised with Histone H2B fused with a tdTomato fluorescent reporter. Arrows indicate mitotic cells. Scale bars: 10µm. **C.** Diagram showing the CRISPR/Cas9 targeting strategy employed to generate the Tet1^EGFP^ knock-in cell line. Mouse embryonic stem cells were transfected with Cas9, a sgRNA targeting Tet1 STOP codon and a repair template for homologous recombination. Coding sequence is indicated in blue, non-coding sequence (3’UTR) is indicated in grey. **D.** Agarose gels showing results from PCR genotyping in homozygous Tet1^EGFP^ mouse embryonic stem cell clones. The location of primers pairs and size of PCR products is indicated in panel C. Homozygous clones C2 and C4 (highlighted in red) were used for live-cell imaging experiments.

**Supplementary Figure 2 (related to Figure 2).**
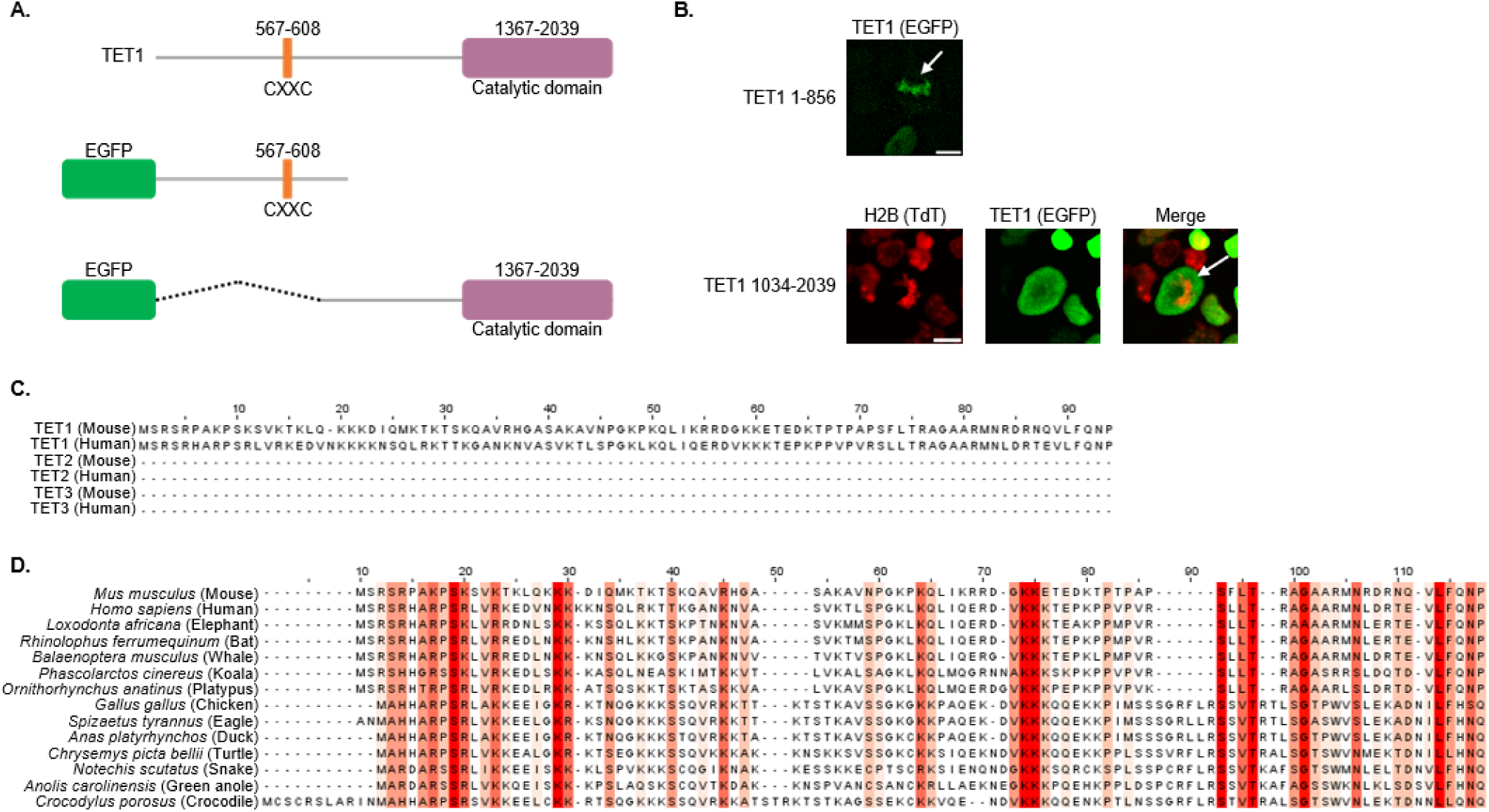
A. Diagram showing TET1 protein structure and different truncations cloned into expression plasmids (EGFP fluorescent reporter) to map the region responsible for mitotic chromatin binding. **B.** Live-cell imaging showing the sub-nuclear localisation of the indicated EGFP-TET1 constructs in mouse embryonic stem cells. Arrows indicate mitotic cells. Scale bars: 10µm. **C.** Protein alignment showing the absence of a region homologous to TET1 N-terminal region (NT) in TET2 and TET3 paralogues in both mouse and human. **D.** Protein alignment showing the conservation TET1 N-terminal region (NT) across amniotes (mammals, reptiles, birds). Strictly conserved residues are highlighted in dark red and conservative substitutions in light red.

**Supplementary Figure 3 (related to Figure 4).**
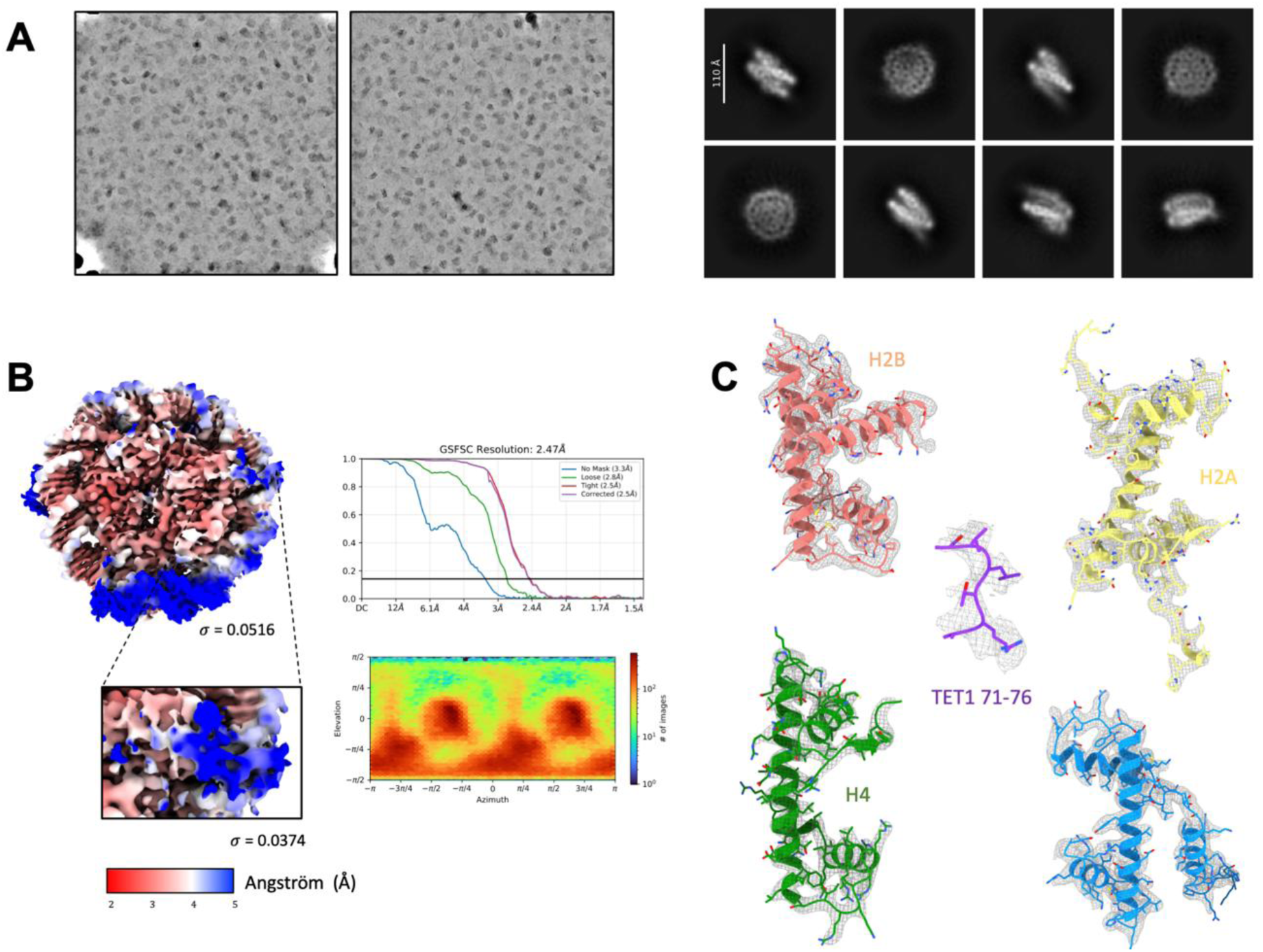
A. Representative denoised micrographs used for particle picking in cryoSPARC (left). Initial particle picking was performed using the Blob Picker and 2D class averages (right) of the TET1(1-93) – NCP complex. **B.** Cryo-EM map coloured according to local resolution from highest resolution (red) to lowest resolution (dark blue). The inset displays TET1(1-93) binding region at different threshold. On the right are shown the gold standard Fourier shell correlation (GSFSC) curves (top) and assigned direction distribution (bottom) of the TET1(1-93) – NCP complex. The GSFSC curves show overall nominal resolutions of 2.47 Å. The angular distribution shown corresponds to the final refinement performed with C2 symmetry. Initial processing and refinement were performed in C1. **C.** The cryo-EM map around the histones and TET1(1-93) is shown in mesh in two views. The side chains of the abovementioned residues are well covered by the map.

**Supplementary Figure 4 (related to Figure 4).**
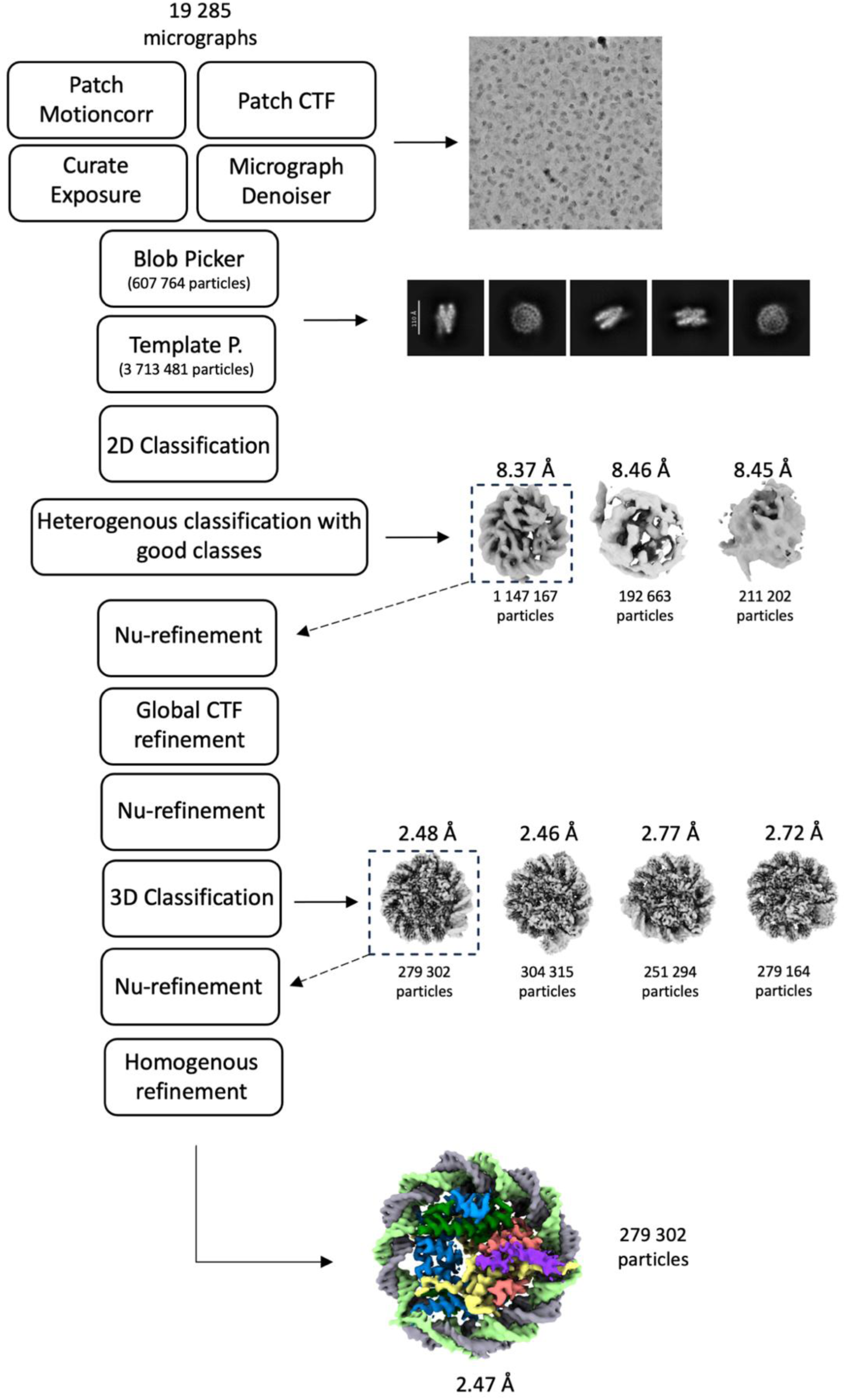
Overview of the cryo-EM processing steps for TET1 in nucleosome bound state. The particles were processed using the software Cryosparc.

**Supplementary Figure 5 (related to Figure 5).**
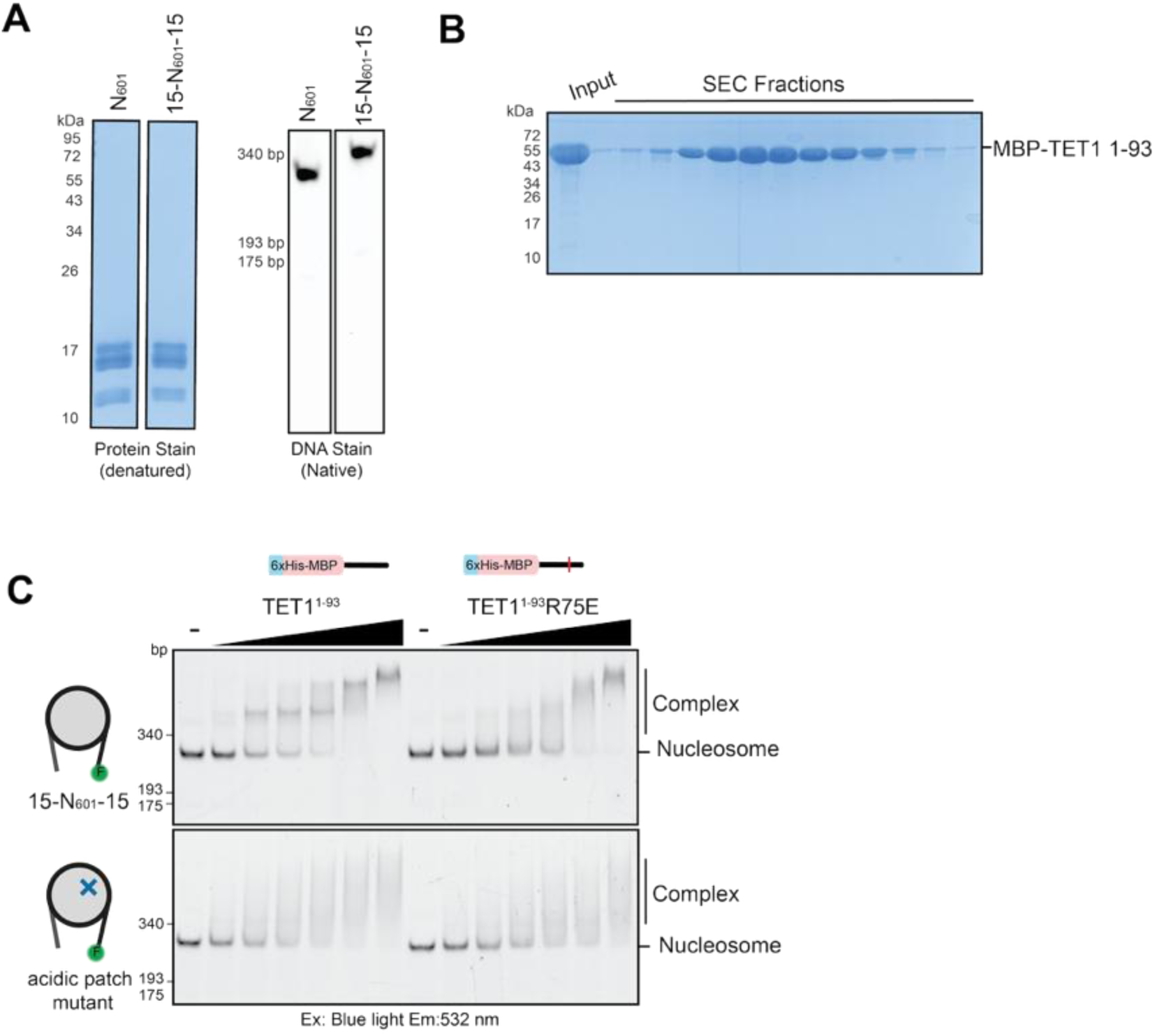
**A**. Representative denaturing SDS-PAGE gel (left) and native polyacrylamide gel (right) of nucleosome core particles (N601) or nucleosomes containing linker DNA (15-N601-15) used for biochemical experiments. **B.** SDS-PAGE gel of final step of 6xHis-MBP-TET1(1-93) purification by size exclusion chromatography (SEC). **C**. EMSA comparing binding of TET1 N-terminal region (1-93) wild-type and R75E mutant to unmodified or acidic patch mutant (H2A^E61A/E91A/E92A^ and H2B^E105A^) nucleosomes wrapped with 5’ FAM (F) labelled 175bp Widom601 DNA. Limiting amounts (2.3 nM) of nucleosomes were incubated with increasing concentrations (0-500 nM) of TET1. Complexes were resolved by native-PAGE and imaged using blue light excitation and 532nm emission filters.

## Other supplementary material

*– Supplementary Video. Timecourse live-cell imaging of endogenous TET1 fused with EGFP during mitosis in mouse embryonic stem cells. Scale bar: 10µm*.

